# Decoding Sequence-Structure-Function-Evolution of basic Leucine Zippers of Aureochromes from Heterokont Algae

**DOI:** 10.1101/2022.05.19.492614

**Authors:** Madhurima Khamaru, Debarshi Bose, Anwesha Deb, Devrani Mitra

## Abstract

The blue light photoreceptor cum transcription factors, Aureochromes (Aureos), are present exclusively in photosynthetic stramenopiles. Co-existence of Light-Oxygen-Voltage (LOV) and basic leucine zipper (bZIP) is unique to Aureos – therefore ideal to study light-dependent DNA binding/transcriptional regulation. Further, Aureos’ inverse effector-sensor topology, resembling several sensory eukaryotic transcription factors, makes them prototypical optogenetic scaffolds. In absence of 3D data, this study aims for a thorough investigation of the bZIP domains from Aureos and others, and their interaction with substrate DNA using tools from sequence/structural bioinformatics, network theory, molecular dynamics simulation and *in vitro* experiments. An in-depth comparison of 173 Aureo/plant/opisthokont bZIPs reveals Aureos’ uniqueness and evolutionary significance in DNA binding specificity as well as dimer stability. An all-atom network analysis on representative bZIP-DNA co-crystal structures, especially the measurement of eigenvector centrality, further adds importance to hydrophobic interactions in the zipper region to stabilize bZIP dimer and facilitate DNA binding in Aureos and other bZIPs. Perhaps the most notable finding is the unique histidine substitution at the basic region of Aureos unlike any other bZIPs. Not only is this residue important for DNA binding, this can serve as a potential switch point in Aureo/bZIP evolution.

**Highlights:**

- Aureochrome is perhaps the only light-responsive transcription factor in bZIP superfamily.
- We draw a comparative between aureochromes and other bZIPs via *in-silico*/*in-vitro* methods.
- Aureochromes form a distinctly separate lineage, midway in bZIP evolution.
- The unique histidine facilitates aureochromes’ interaction with cognate DNA substrates.

## 1. Introduction

Regulation of gene expression is essential for the survival of every life form. Transcription factors (TFs), upon binding with the cognate DNA, turn genes on/off at precise spatio-temporal resolution. Different classes of transcription factors exist. They differ on the basis of their DNA binding structural motifs and activity. Examples include basic leucine zipper (bZIP), basic helix-loop-helix (bHLH), Zinc-coordinating DNA-binding motifs (Zinc finger), helix-turn-helix (HTH), beta scaffold factors like STAT (Signal transducer and activator of transcription proteins), MADS (MCM1, AGAMOUS, DEFICIENS, and SRF) box, HMG box (high mobility group box), TBP (TATA-binding protein), Beta barrel alpha helix etc. Among these, bZIPs comprise one of the largest group of eukaryotic transcription factors (Ji et al., 2018). The uniqueness of bZIPs lies in their conserved nature of structure and sequence specificity. They possess N-X7-R/K signature motif and binds mostly to the *cis* acting DNA elements of a promoter having an ‘ACGT’ core sequence (Wang et al., 2018) (Rodríguez-Martínez et al., 2017). Interdigitation between the partner helices leads to the dimerization of bZIP — resulting in a superimposed coiled coil structure or ‘zipper’ (Bader & Vogt, 2006)(Sinden, 1994). This zipper-like structure is extended by the N-terminal helical segment, which can bind to the DNA major groove via basic amino acid residues(Glover & Harrison, 1995). Dimerization is essential for the recognition of substrate DNA and transcriptional regulation. The C-terminal zipper motif consists of heptads, which can be represented by a helical wheel. The residues are conventionally marked from ‘a’ to ‘g’ (Grigoryan & Keating, 2006), where leucines generally occupy every ‘d’ position, whereas the hydrophobic residues including leucine/isoleucine dominate ‘a’ (Bader & Vogt, 2006).

However, presence of a conserved asparagine at ‘a’ position in the middle of zipper could result in destabilization of the oligomer (Harbury et al., 1993). The ‘b’, ‘c’ and ‘f’ positions are generally filled by polar residues. The charged residues at the ‘e’ and ‘g’ positions also aid in the zipper dimer stability (Deppmann et al., 2004)(Podust et al., 2001). Leucine residues, owing to their hydrophobic side chains, create ‘knobs’ which can fit into the ‘holes’ generated in between the side chains of amino acids present in adjacent helices (Bader & Vogt, 2006). At the other end, the basic amino acid residues complement well with the negatively charged phosphate backbone of DNA — this results in favorable electrostatic interactions. Besides, they can also participate in hydrogen bonding/polar interactions and can even engage in non-polar interactions with the portions of extended side chains. Besides phosphates, the nitrogenous base and deoxyribose sugar are also involved in hydrogen bonding, van der Waal’s as well as stacking interactions. The affinity of bZIP TFs towards the cognate target DNA sequences is imparted by the sequence composition of its basic region as well as the length of the leucine zipper (Hakoshima, 2005a). The dimerization partner is also crucial for its sequence specific DNA binding (Pogenberg et al., 2014). Their ability to bring in change in the binding specificity upon hetero-dimerization is indicative of this (Busch & Sassone-Corsi, 1990). The electrostatic attraction and repulsion forces, acting between the polar residues that flank the hydrophobic interaction surface of the zippers, are major deciding factors behind the formation of homo/heterodimers (C. R. Vinson et al., 1993). The oppositely charged residues located at ‘g’ and the following e’ positions across the hydrophobic interface, form the inter-helical ion bridges. The interactions between d-d’, a-a’, g-e’ are vital for the zipper formation and stabilization (Podust et al., 2001).

Widespread among the regulatory proteins of gene expression, the bZIPs play a crucial role in cellular proliferation and differentiation in eukaryotes. It is known to be the ancient DNA binding module found across three kingdoms of life (Hakoshima, 2005b). Several groups of opisthokont and plant bZIPs are well defined and characterized. Few homologs of bZIPs are also evident in fungi, algae, protists and even in bacteria. However, the only blue light (BL) responsive photoreceptor with a bZIP TF activity has been recently discovered (Takahashi et al., 2007) in Aureos – found exclusively in photosynthetic stramenopiles (Rayko et al., 2010). The flavin binding Light-Oxygen-Voltage (LOV) BL sensor at the C-terminus, makes Aureos different from any other conventional bZIPs. Aureos perform organism specific diverse functionalities [**SI-1**] e.g. - branching in *Vaucheria sp* (Takahashi et al., 2007), cell cycle progression in *Phaeodactylum sp* (Kroth et al., 2017), lipid accumulation in *Nannochloropsis sp.* (Huang et al., 2014) etc. The broad spectrum functions of Aureos include promotion of high light acclimation, photo protection in diatoms etc (Huysman et al., 2013). Considering maximum penetration of BL in sea water, most of the physiological and developmental responses are BL dependent in photosynthetic marine algae. However, it lacks phototropins (Rayko et al., 2010), which is one of the major BL photoreceptors in land plants. Aureos instead aid in the optimization of photosynthesis (Mann et al., 2020) under varied light conditions. In fact, Aureos’ significance in carrying out BL-mediated gene expression is not only important to study BL signaling in photosynthetic marine algae, they are potentially relevant for optogenetics as well. The direct DNA binding ability of Aureo in response to BL without an intricate signaling mechanism makes Aureo an ideal optogenetic scaffold. They are capable of bringing in faster changes in gene expression under fluctuating light conditions (Mann et al., 2020). Dual presence of a LOV photoreceptor and a bZIP TF domain empowers Aureo to serve as a natural optogenetic tool (Hepp, Trauth, Hasenjäger, et al., 2020) (Duan et al., 2018) (Jerng et al., 2021). In contrary to commonly-sighted domain topology in most LOV photoreceptors, the inverse effector-sensor arrangement in all Aureo is similar to that in several eukaryotic TFs (Matiiv & Chekunova, 2018) — a feature suitable for developing synthetic sensory TFs (Boral et al., 2022) as well as optogenetic designs (S. Banerjee & Mitra, 2020). While a substantial amount of literature is available against Aureo LOV sensors especially on their sequence-structure-function (Mitra et al., 2012) (A. Banerjee, Herman, Kottke, et al., 2016) (Hepp, Trauth, Hasenj, et al., 2020) (Heintz & Schlichting, 2016), structure based mechanistic insights is lacking for the effector bZIPs. And till date, no experimental structure corresponding to full-length or even the bZIP-linker-LOV module of Aureo is available in PDB. Therefore, a comprehensive sequence-structure characterization of this effector bZIP is attempted using several *in-silico* tools e.g. for phylogenetic studies, network theory, modelling, docking and molecular dynamics (MD) simulation to understand the formers’ uniqueness and significance. We include all available varieties of Aureo_bZIPs and compare them with other traditional bZIP TFs found across plants and opisthokonts to understand the uniqueness and evolutionary significance. We further validate our *in-silico* results and assess the DNA binding ability of Aureo_bZIPs using *in-vitro* electrophoretic mobility shift assay (EMSA) after cloning, overexpression and protein purification. In this study, we attempt to draw an effective comparative between the bZIP domains of Aureos and the rest *via* a holistic sequence-structure-function characterization.

## 2. Materials and Methods

### 2.1. Retrieval of bZIP sequences from different organisms

The amino acid sequences of Aureos and other representative bZIP proteins from plants and opisthokonts were collected from the UniProt (Apweiler et al., 2018) protein sequence database. BLASTP (Altschul et al., 1990) search was also performed against the database of non-redundant protein sequences using available bZIP sequences as queries. A few sequences, especially those of Aureos from *Phaeodactylum sp*. were obtained from the available literature (Costa et al., 2013). Amino acid sequences of the 78 bZIP proteins from plants, 69 bZIPs from opisthokont species and 26 Aureo bZIP sequences were shortlisted for further analyses. The respective domain boundaries of the bZIPs were finalized either from ‘conserved domain’ analysis of NCBI and Pfam, or from PSI-BLAST or domain BLAST of NCBI.

### 2.2 Sequence comparison and molecular evolutionary analysis

A total of 173 bZIP sequences thus retrieved, were subjected to multiple sequence alignment using Clustal omega (Sievers et al., 2011) algorithm — first group-wise and then collectively. This gave us a preliminary understanding on the subtle differences in bZIP sequences that could lead to differences in DNA binding activity. The sequences were further considered for the construction of a phylogenetic tree using MEGA X programme (Kumar et al., 2018). The corresponding accession numbers of all the database entries of these bZIP proteins along with the names and source organisms are listed in the table (**SI-2**). For molecular evolutionary analysis, all 173 bZIP sequences were first aligned using MUSCLE algorithm, inbuilt in MEGA X. The most rational tree of all the selected bZIP proteins was drawn using the neighbor joining (NJ) method, which is simple and fast. Poisson model, which considers equality of substitution rates among all the sites and assumes equal frequencies of amino acids at the time of correction of the multiple substitutions at the same sites, was followed as the substitution method. The phylogeny was tested through bootstrap analysis where the number of performances of bootstrap replications for each node was kept at 1000. The redundant and partial sequences were discarded. Only the reviewed and annotated sequences were considered. The guide tree was then represented and labeled by the online phylogenetic tree manipulation tool iTOLv5. The clades in red and green represent the maximum and minimum values of bootstrap respectively. A covariance analysis on Aureo bZIPs were further performed using the GREMLIN (Kamisetty et al., 2013) server. Default parameters like 4 iterations, >75% query coverage and 10^-10^ /10^-4^ E-value cut-off were chosen as appropriate for this study.

### 2.3 Residue Interaction Graphs to understand bZIP-DNA interactions

The protein-DNA complex structure can be represented as residue interaction graph (RIG) or an all-atom network, where nodes (amino acids/nucleotide residues) are linked by edges within a given distance cut off (e.g. 4.5 Å). To investigate further on the bZIP-DNA complexes, RIGs were first generated from eight representative homo/heterodimeric bZIP co-crystal structures from all the bZIP subfamilies, using the Biographs module of Biopython (Cock et al., 2009) and NetworkX (Hagberg et al., 2008). Eight structures representing different bZIP subgroups are - 2WTY (MAF), 1JNM (JUN-CRE complex), 1GD2 (PAP), 1YSA (GCN4), 1FOS (JUN-FOS heterodimer), 1NWQ (CEBP), 1DH3 (CREB) and 2H7H (JUN-AP1 complex). The RIGs were generated considering all atom-pair contacts within a given distance of 4.5 Å. Centrality measures as simple as degree centrality, geodesic/shortest path based closeness and betweenness centrality have been extremely helpful to identify influential nodes from residue-level interaction data (Deb et al., 2020). However, to capture all communication at the residue level, we calculated eigenvector centrality (EC) of the constructed RIGs. How well connected a node is, in terms of its connectivity with other well-connected nodes of a network, can be measured by EC. The weighted sum of the centralities of all nodes, which are connected to node *‘i’* by an edge *‘A_ij_’*, is defined as EC of node *‘i’* (*C_i_*) (Negre et al., 2018)- *C_i_* = *ε*^-1^∑*_j_*_=1_*^n^A_ij_c_i_*, where ‘*c*’ denotes the eigenvector associated with eigenvalue *ε* of ‘*A*’.

The distribution of EC was plotted next. We studied the value of Z (X) = (X-<X>)/σ, where <X> and σ represent mean and standard deviation respectively. For subsequent analysis, we therefore considered only those values of X which are associated with Z (X) > 1.5.

### 2.4 Modeling and docking of Aureo bZIPs - comparison with other bZIPs

Till date, no experimental 3D structure corresponding to the bZIP region of Aureo is available in RCSB PDB. Hence, the bZIP sequences, representatives from all five subtypes of Aureos sourced from *Ectocarpus/Saccharina* sp, were individually modeled using SWISS-MODEL (Waterhouse et al., 2018). The best models, selected on the basis of best Ramachandran parameters, were next energy minimized using Chimera (Pettersen et al., 2004). The energy minimized structures were next subjected to molecular docking analysis using NPDock (Tuszynska et al., 2015). Review of literature suggests that the bZIP domain of Aureo binds to a consensus DNA sequence TGACGT, named ‘Aureo-box’(Matiiv & Chekunova, 2018) (Takahashi et al., 2007). So, for the *in silico* docking of modeled Aureo_bZIPs with DNA, ‘Aureo-box’(TGACGT) containing CRE (cAMP responsive element) DNA (A. Banerjee, Herman, Kottke, et al., 2016) from co-crystal structure of CREB bZIP-CRE DNA [(PDB ID: 1DH3, (Schumacher et al., 2000) was used as the DNA substrate /ligand. Preliminary knowledge on protein-DNA interaction interface, as acquired from sequence-structure superposition with solved bZIP-DNA co-crystal structures and residue interaction graphs was used as input in docking. The energy parameters, interface interactions as well as buried surface area on the best scoring structures of Aureo_bZIP-DNA complexes were calculated using PDBePISA (Evgeny, 2010). In order to perform a detailed comparative study between aureochromes and the rest bZIPs, the hydrogen bonding and polar interactions at the basic region of bZIP-DNA interfaces of all were analyzed using Chimera (Pettersen et al., 2004) and Pymol (Delano, 2002). *In silico* mutagenesis was performed in few docked structures to understand the functional as well as evolutionary significance of N-terminal basic region residues of Aureo_bZIPs.

### 2.5 System Setup, Molecular Dynamics Simulations and Post-Simulation Methodologies

Following phylogeny and docking analyses, unique placement of histidine was noticed exclusively at the basic region of Aureos and not in others. In order to investigate the specific contribution of this residue in DNA binding activity, Molecular Dynamics (MD) Simulations were performed using the GROMACS (version 2020.1) (Pronk et al., 2013) (Van Der Spoel et al., 2005). The docked structure of *Ectocarpus siliculosus* Aureo1_bZIP-DNA complex (*Es*Aureo1_wild) and its *in-silico* mutant versions (*Es*Aureo1_HA and *Es*Aureo1_HS) were considered as initial files for MD simulation. AMBER99SB-ILDN force field was applied. The docked bZIP-DNA complex was solvated using TIP3P water molecules and was limited into a cubic lattice of dimensions 1.2 x 1.2 x 1.2 nm^3^. The structures had positive charges at normal physiological pH conditions, thus the addition of chloride (Cl^-^) ions ensured the simulation systems’ electrical neutrality. The entire molecular structure was then relaxed by performing energy minimization, starting with the steepest descent and followed by the conjugate gradient algorithm. To remove the solute-solute and solute-solvent steric clashes and allow the system to adapt to its environment stably, a harmonic restraint was imposed on the solute particles keeping the solvent particles unrestrained, for over 50000 energy minimisation steps for both algorithms. Before moving forward, the potential energy (E_pot_) of the system was checked to be a negative value, since it is correlated with the sufficient stability of the system required for simulation. The equilibration process was conducted in two steps. First, the isothermal-isochoric or NVT (constant number of particles-volume-temperature) ensemble, followed by an isothermal-isobaric or NPT (constant number of particles-pressure-temperature) ensemble was conducted for 100 ps each, with position restraining for the non-water residues and constraints of the H-bonds. A leapfrog integrator was used to analyze the dynamic system numerically. The Berendsen thermostat algorithm was used for re-scaling the particle velocities in the simulation model to relax the system and regulate the temperature inside the box to be at around 300 K. The Particle Mesh Ewald (PME) method was utilized for rapid stochastic relaxation of the system to equilibrium and to calculate long-range electrostatics. The pressure fluctuation of the simulation system was accounted for by running an average over the entire length, which was compared with the reference pressure set as 1 bar and an isothermal compressibility allowance of 4.5 x 10-5 bar^-1^ for water. The LINCS (LINear Constraint Solver) algorithm was used for bond (involving hydrogens) restraining. An integration time step of 2 fs was applied and the log file/ coordinate update was done every 1 ps. The system, thus prepared, was finally exposed to the subsequent forces and ran for 60 ns (20 ns at a stretch for 3 consecutive times), at 300 K temperature.

The data obtained post-simulation (as .xvg files) were plotted using the XMGRACE plotting tool. PyMOL (Delano, 2002) software was used for visual assessment of the bZIP-DNA complexes during the simulatory period. The GROMACS in-built analysis package (Van Der Spoel et al., 2005) was extensively utilized to elucidate all the analytical data for 60 ns MD simulation run, which were compared for the wild-type and the mutant variants.

### 2.6 Cloning of wild type Aureochrome-1 from Ectocarpus siliculosus

The pUC-57 plasmids harboring the codon optimized synthetic gene corresponding to *Ectocarpus siliculosus* aureochrome1 (*Es*Aureo1; Uniprot accession no.-D7FMH8) was purchased. PCR amplification of the bZIP-linker-LOV sequence of wild type *Es*Aureo1 (*Es*Aureo1_Wild) DNA sequence was carried out using the following forward primer (FP) and reverse primer (RP) respectively-5’CTAGGCTAGCATGAAAGATCTGACCGAAGAACAGCGCATTG3’; 5’AGCTGAGCTCTTATTTGCTCGGGCCCGCCAGCTG3’. The *Es*Aureo1_wild was then cloned into the pET28a expression vector at the NheI/SacI restriction sites.

### 2.7 Site directed mutagenesis

To uncover the role of unique histidine substitution at the basic region of Aureos, His362 residue in *Es*Aureo1_wild was next mutated to alanine and serine. These mutations led to the formation of *Es*AureoHA and *Es*AureoHS respectively. Following primers were used to generate *Es*AureoHA and *Es*AureoHS constructs respectively-*Es*AureoHA_FP:5’CGCAACCGCGAAGCGGCGAAACGCAGC3’, *Es*AureoHA_RP:5’GC TGCGTTTCGCCGCTTCGCGGTTGCG3’;*Es*AureoHS_FP:5’CGCAACCGCGAAAGCGCGAAACGCAGC3’; *Es*AureoHS_RP:5’GCTGCGTTTCGCGCTTTCGCGGTTGCG3’.

### 2.8 Overexpression and purification of WT and mutant proteins

*Es*Aureo1_wild and its mutatants - *Es*AureoHA and *Es*AureoHS were transformed into the *Escherichia coli* C43 strain to express the wild type protein along with its mutants. Colonies obtained through transformation were then inoculated into the kanamycin containing Luria-Bertani (LB) (*HiMedia*) broth cultures and grown overnight at 37°C. In each case, 1% inoculum was transferred to the fresh media and grown at 37°C till OD_600_ reached 0.6. Following IPTG (*Promega*) induction, the cells were grown in the dark at 21°C overnight. Bacterial cells were then harvested by centrifugation at 8,000 rpm for 10 minutes. The cell pellet was then dissolved in lysis buffer (20 mM Tris, pH 8; 10 mM NaCl; 10% Glycerol) and incubated with protease inhibitor cocktail (*Sigma*), a pinch of lysozyme (*SRL*) and phenylmethylsulfonyl fluoride (PMSF, *Promega*). The dissolved cell pellet was then subjected to ultrasonic vibrations (*Hielscher*) and high-speed centrifugation (*Thermo Fisher Scientific*) at 12500 rpm for 1 hour at 4°C. Following centrifugation, the supernatant was collected and subjected to Ni-NTA affinity chromatography (Qiagen). Following loading and wash, hexa-histidine tagged protein fractions were eluted using the elution buffer (20 mM Tris, pH 8; 10 mM NaCl; 100 mM Imidazole; 10% Glycerol). The eluted fractions containing the desired protein were then subjected to PD-10 desalting column (*Sigma*) for the removal of excess imidazole. The desalted protein was then concentrated using Amicon® Ultra Centrifugal Filter. The quality of protein was checked using SDS-PAGE (*Bio-Rad*). Further purification by Heparin/gel filtration chromatography resulted in extremely poor yield. Therefore, affinity purified and desalted protein fractions were used for EMSA studies, carried out at least in triplicates from multiple batches of protein purification. The concentration of the wild type as well as the mutant proteins was specifically determined with respect to flavin [at 450 nm, considering the extinction co-efficient of flavin (12500 M^-1^cm^-1^) using UV-Vis spectroscopy (*Shimadzu*)].

### 2.9 Electrophoretic Mobility Shift Assay

HPLC purified and lyophilized single7stranded oligonucleotides of 24bp containing ‘Aureo-box’ sequence, ‘TGACGT’, (FP: 5’TGTCCTTGGCTGACGTCAGCCAAGCAC3’;RP:5’GTGCTTGGCTGACGTCAGCCA AGGACA3’) were purchased (*IDT*). Following resuspension in annealing buffer (10 mM Tris, pH 8.0; 20 mM NaCl) the complementary DNA strands in equimolar amounts were annealed by heating at 95°C for 2 mins followed by gradual overnight cooling. The binding buffer (50 mM Tris, pH 8.0; 50 mM NaCl; 1.25 mM MgCl_2_; 0.01 mg/mL BSA; 20% glycerol) was next added to the double stranded DNA (dsDNA) to achieve a final concentration of 0.5 uM. Each protein was then serially diluted and added to the solution of dsDNA and binding buffer mixture. A total of nine samples were prepared for protein-DNA mixtures with varying protein concentrations. The substrate DNA minus protein was considered as negative control. All samples including the control were incubated on ice under BL for 45 minutes prior to the commencement of electrophoresis. The electrophoretic run was performed using Bio-Rad’s Mini-PROTEAN system at 4°C under BL-illuminated condition. The samples were resolved at 150 Volts at 4°C for one hour. Upon the completion of electrophoresis, the native gel was stained with SYBR Gold nucleic acid stain (*Thermo Fisher Scientific*) for 45 minutes and imaged in the Bio-Rad ChemiDoc MP system.

## 3. Results

### 3.1. Sequence retrieval and analysis

A total number of 173 bZIP sequences inclusive of 78 from plants, 69 from opisthokonts and 26 from Aureos are considered [**SI-2**] herein to assess amino acid sequence composition/conservation/variations. The conservation of leucines/non-polar amino acid residues at every a/d position in the C-terminal dimerization domain/zipper region of Aureos corroborate well with other bZIPs. However, we do observe few remarkable variations at the N-terminal basic region of Aureo_bZIPs when compared with others. The basic region responsible for DNA recognition possesses a signature sequence of nine amino acid residues [N-(X)_7_-K/R] [**SI-3**]. A histidine is uniquely found at the 4th position [N-(X)_2_-**H**-(X)_4_-R] of the basic region in Aureos [**Figure-1.a**] unlike others. While in plant-bZIPs serine is the invariable occupant (except plant bZIP subfamily D, F) [**SI-3.**]; alanine is mostly present in opistokont bZIPs, excluding CREB, CREB3, ATF6 [**SI-3**]. Therefore, the basic region sequence of opisthokont and plant bZIPs can be broadly designated as [N-(X)_2_-**A**-(X)_4_-K/R] and [N-(X)_2_-**S**-(X)_4_-R] respectively. Notably in Aureo2, histidine is substituted with leucine [**Figure-1.a**]. An in-depth investigation into all the Aureo further divulged that *Sj*Aureo2 despite being designated as a type-2 Aureo, possesses histidine [**SI-3**]. This observation can be justified from the phyletic pattern [**Figure-1.b**], where *Sj*Aureo2 clubs with other Aureo paralogs rather than the Aureo2 cluster. As neither histidine nor leucine is found at the debatable 4^th^ position in any other bZIP across three kingdoms of life, their persistence in aureos could bear an evolutionary significance. Again, the presence of threonine as the 8^th^ residue in Aureo2 basic region [N-(X)_2_-LA-(X)_2_-**T**R] [**Figure-1.a**] is also notable considering the presence of serine at the homologous position in most other Aureo bZIPs. Incidentally, the 8th position of all bZIPs is occupied mostly by serine or cysteine. A covariation of histidine at 4th and serine at the 8th position is further detected in the co-variation analysis using GREMLIN (Kamisetty et al., 2013)[**Figure-1.c**]. This probably explains why both histidine and serine are simultaneously replaced by leucine and threonine respectively in Aureo-2 paralogs.

**Figure-1:**
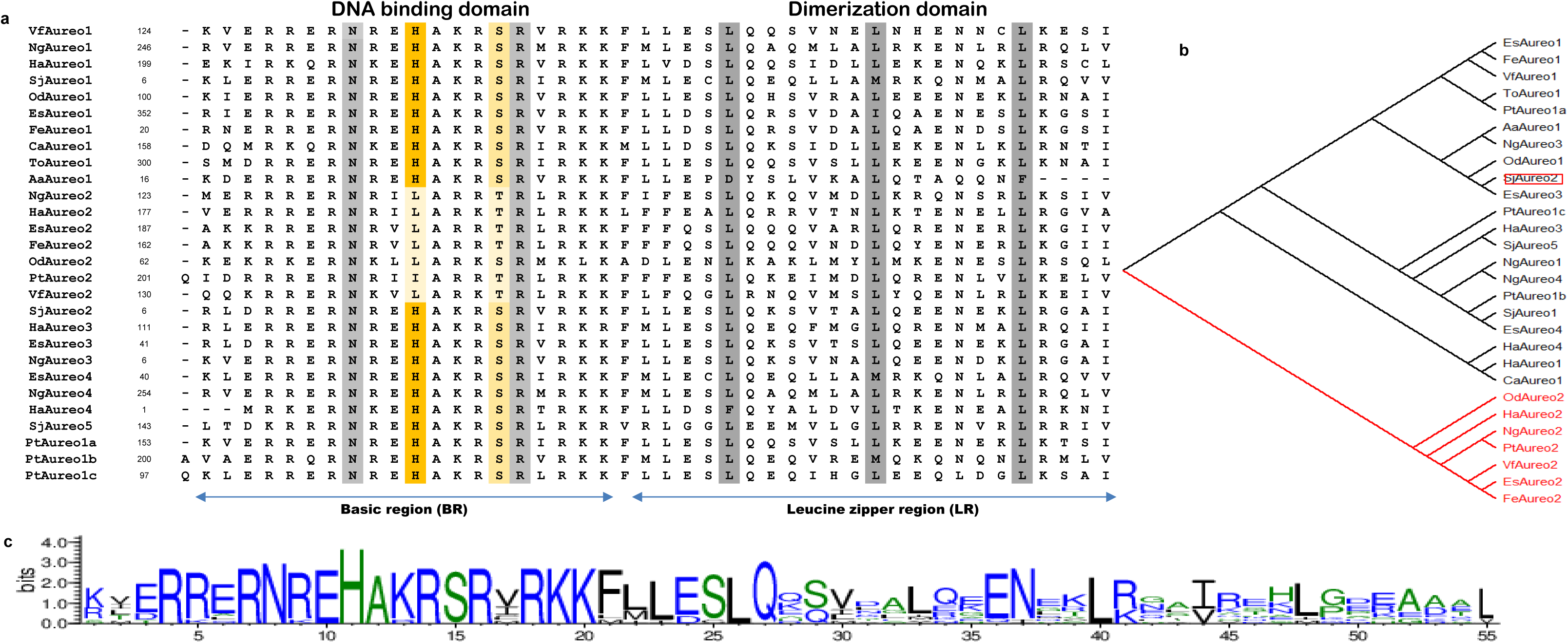
a. Multiple sequence alignment of the bZIP domains of Aureos b. Phylogenetic tree comprising all the aureochrome homologs. c. Weblogo result of the co-evolution study of aureochrome1 at E-value cut-off 10^-10^.

### 3.2 Phylogeny estimation and course of bZIP evolution

The evolutionary position and the phylogenetic relationship of all selected 173 bZIP proteins from plants, opisthokonts and Aureos can be inferred from the phylogenetic tree construction [**Figure-2**]. However, the deeply rooted divergence within the ancient nodes may not have been detected due to the comparatively shorter length of bZIP domains. Nevertheless, three distinct lineages from a common ancestor have been observed. The first lineage includes BATF, PAR, and E4BP4. The second lineage comprises opisthokont bZIP families such as CREB2, ATF2, CEBP, ATF3, FRA, and FOS along with the viral bZIP family Zta. The remaining plant and opisthokont bZIPs are found to cluster together with Aureo bZIPs in the third lineage. However, the last common ancestor of the third lineage further diverges into two distinct clades. One is solely represented by all Aureo_bZIPs, while the other splits further into two branches: the opisthokont bZIPs (e.g., PAP, GCN4, JUN, MAF, CREB3, CREB, and ATF6) and all plant bZIPs (e.g., A, B, C, D, E, F, G, H, I, and S). However, the most appropriate sister clade or immediate ancestor of Aureo bZIPs could not be ascertained. The phylogenetic tree of all bZIPs illustrates the tendency of Aureo_bZIPs to club together in the same clade, forming a distinct sub-lineage without intermingling with bZIPs either from plants or opisthokonts. Most Aureo paralogs within the Aureo-bZIP clade align by subtype, with a few exceptions, while the remaining ones lose their distinctiveness and cluster randomly [**Figure-2**]. In summary, the phylogeny estimation of bZIPs from multiple genomes not only identifies the ancestor-descendant relationships among different bZIP members but also highlights the evolutionary relationship of Aureo_bZIPs with the rest.

**Figure-2:**
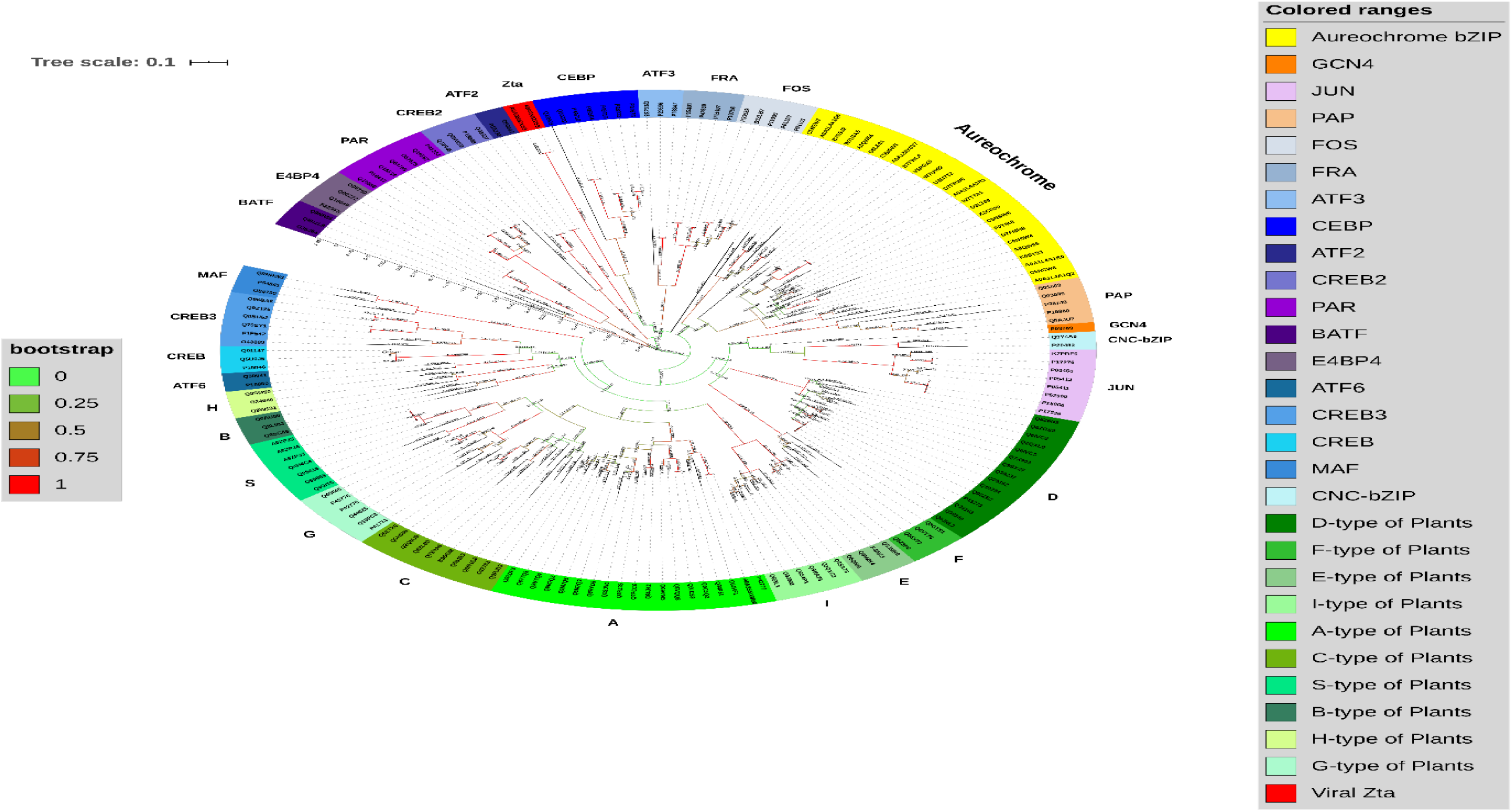
Phylogenetic tree of the bZIPs from plants, opisthokonts and Aureos. Different subgroups of bZIPs are represented by different colours. Highest to lowest bootstrap values are colour coded as green and red respectively.

### 3.3. Network analysis and bZIP dimer stability towards substrate DNA interaction

Aureo bZIPs are unique in the sense that they are BL-regulated. In fact, the LOV → bZIP signaling relies on an allosteric pathway (A. Banerjee, Herman, Serif, et al., 2016). The resulting conformational changes at the bZIP-substrate DNA interface further promotes DNA binding without much alteration in DNA binding affinity. In absence of full-length experimental structure of any Aureo at dark and light states, it would be difficult to precisely identify the intervening residues that transduce light energy into DNA binding. Despite significant homology observed between bZIP and LOV domains individually with other bZIP/LOV members respectively, the existence of heterogeneous linkers with varying sequence composition and length further complicates this problem. While bZIP dimerization is an essential pre-requisite of DNA binding activity, Aureos contain the only bZIPs that are connected with a BL-sensor. Therefore, one crucial question is: how does an Aureo bZIP compare with other bZIPs? We therefore constructed all-atom networks from eight co-crystallized bZIP-DNA complexes. While centrality measures are frequently used for network analysis, in this study, we particularly focused on Eigenvector Centrality (EC) – especially given their implications in allostery (Negre et al., 2018). EC that considers both the number of connections made by a node as well as the flow of information associated with them, EC has been successfully used to identify residues responsible for allosteric communication in enzymes (Negre et al., 2018). We did not consider Aureo model structures for RIG analysis because structural artifacts could affect the result. We therefore calculated EC from networks constructed upon eight co-crystal structures of homo/heterodimeric bZIP-DNA complexes [PDB IDs: 1DH3, 1FOS, 1GD2, 1JNM, 1NWQ, 1YSA, 2H7H and 2WTY] [**SI-4**]. Residues with high EC values with Z(X) > 1.5 are only considered and marked in green [**Figure-3**]. Additionally, two to three residues from monomeric units showing highest EC values are color coded in red to check any further significance. The residues involved in DNA binding are marked in blue [**Figure-3**]. In all cases, residues with high EC values are found to congregate at the zipper region, albeit at variable distances from the basic region. Residues with highest EC values specifically occupy ‘a’ and ‘d’ positions of a heptad in every single case, as seen in the figure as well as multiple sequence alignment [**Figure-3.b**]. We find this result extremely interesting considering well-known hydrophobic interaction between residues at positions ‘a’ and ‘d’ towards leucine zipper dimerization, its stability as well as their contribution in allosteric regulation (Cruz et al., 2023). In terms of EC, after ‘a’ and ‘d’, the next important positions happen to be ‘e’ and ‘g’. As emphasized in our earlier study (Khamaru et al., 2024), the electrostatic compatibility in between ‘e’ and ‘g’ can singularly govern bZIP homo/heteromerization(C. Vinson et al., 2002)(Krylov et al., 1994). Sequence comparison reveals that the homologous residues in Aureos corresponding to high EC values in other bZIPs are mostly conserved (similar if not identical) except for the ‘d’ position in the 4th heptad. This particular position in Aureo is occupied by polar or even charged amino acid residues, whereas this is always leucine in other bZIPs. Overall, our EC analysis combined with sequence alignment reveals that Aureo zippers and the rest are generally comparable. However, precise contribution of important heptad residues (a/d/e/g) in allosteric regulations and/or signal transduction, as predicted by EC analysis, is subjected to further experimental validations.

**Figure-3:**
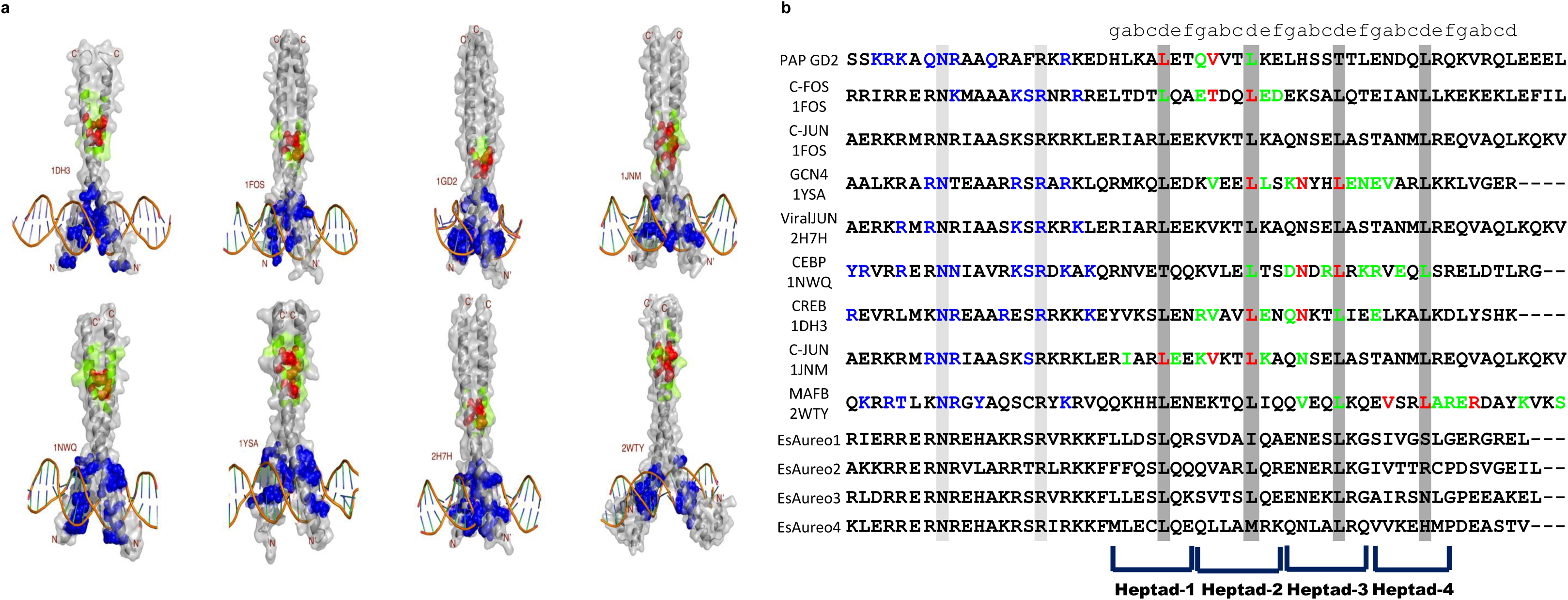
a. DNA-bZIP complex where the residues identified as important from eigenvector centrality measure are colour coded. Green colour denotes EC value higher than 1.5, Red colour indicates knob like structure (d & a) with the highest EC value, blue colour represents DNA binding residues. b. Multiple sequence alignment showing DNA binding residues (coloured blue) and residues with higher EC values (coloured green/red).

### 3.4. Decoding bZIP-DNA interactions - insights from modeled and docked Aureo subtypes

Following *Es*Aureo1, *Es*Aureo2, *Es*Aureo3, *Es*Aureo4 and *Sj*Aureo5 homodimer modeling and energy minimization, each one was subjected to *in silico* docking experiments with the substrate DNA sequence encompassing a TGACGT core which is known to be the binding sequence for Aureo bZIPs, termed as ‘Aureo-box’. The best scoring complex structures were selected and bZIP-DNA interfaces were characterized based on hydrogen bonding/salt bridge/van der Waals’ interactions, interface area and energy. The bZIP-DNA structures of all five Aureo-bZIP representatives are depicted in [**Figure-4**]. The accuracy of our docked structures was further assessed by checking their RMSD when superimposed with the modeled/docked structure obtained through alpha fold. To test the reliability of our docked structures, we conducted PDBePISA (<u>P</u>rotein Interfaces, <u>S</u>urfaces and <u>A</u>ssemblies) analysis and compared them with experimental data on bZIP-DNA complex [1DH3 and 1FOS representing homo- and heterodimer-DNA interactions respectively]. The emphasis was given on three parameters – the interface area, solvation energy and number of non-covalent interactions, precisely hydrogen bonding and salt bridge interactions. While the interface area between monomeric units of protein chains ranges from ∼900 Å^2^-1200 Å^2^ (homodimer to heterodimer respectively), that between protein-DNA ranges from ∼300-400 Å^2^. A negative value of gain in solvation free energy (Δ^i^G) upon interface formation in both protein-protein/DNA complex formations indicate positive affinity between them and the hydrophobic interface. In the case of all homodimeric Aureos [*Es*Aureo1-4, *Sj*Aureo5], the interface area is very similar to homodimeric 1DH3/1JNM, with an average of 915 Å^2^. Aureo-DNA interface area ranges in between ∼350-450 Å^2^ with a very similar number of hydrogen bonding/salt bridge interactions like in experimental structures. Further, the Δ^i^G for both protein-protein/DNA in Aureo-DNA complexes are not only negative but the values are very similar to those of native co-crystal structures. The N-terminal basic regions being the key determinants behind DNA binding specificity of bZIPs, they were first screened for polar including hydrogen bonding interactions with the DNA bases. At first a detailed analysis of the hydrogen bonding interactions involving the basic region and the substrate DNA from experimentally derived co-crystal structures of the bZIP-DNA complexes [**SI-5**] was performed to determine the segment of bZIP crucial for the DNA binding and to identify the crucial residues within the basic region that facilitate DNA binding. Analysis of the bZIP-DNA interaction patterns of all the co-crystal structures revealed multifarious specificities of bZIP proteins despite overall structure-sequence conservation. Since, the five subtypes of Aureo_bZIPs exhibited minor difference in residue composition of the signature motif of bZIP meant for DNA binding, all the five Aureo_bZIPs were docked with DNA [**Figure-4**] and were then screened for the hydrogen bonds and polar interactions present between the basic region residues and the substrate DNA bases to understand whether histidine takes part in the interaction with DNA. In *Es*Aureo1-DNA complex, His362 (NE2) is found to interact with dT’8. *Es*Aureo4 also exhibits interaction of His50 (NE2) with DNA. His153 (NE2) of *Sj*Aureo5 interacts with OP2 of dG’6. All the polar interactions detected at the protein-DNA interface of the docked structures are color coded and presented in tabular format [**SI-6**].

**Figure-4:**
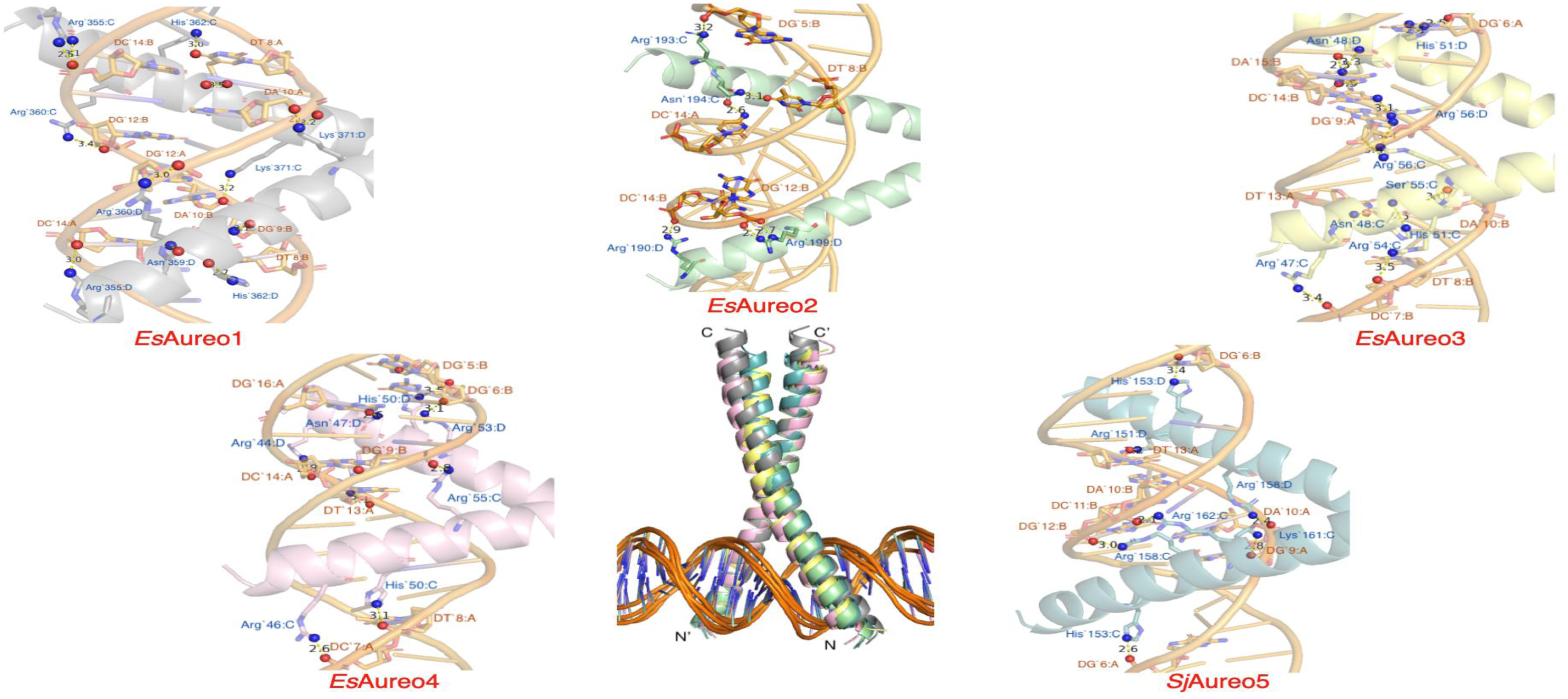
Docked bZIP structures of *Ectocarpus siliculosus* (*Es*) aureochrome1 (gray), aureochrome2 (pale green), aureochrome3 (yellow), aureochrome4 (pink) and *Saccharina japonica* (*Sj*) aureochrome5 (teal) bZIPs in complex with substrate DNA sequences containing ‘Aureo-box’, TGACGT motif. Interactions between amino acid residues of the DNA recognizing the basic region of bZIPs and the corresponding nucleotide of substrate DNA are labeled.

Characteristic of all bZIPs, in Aureo-DNA complexes too, the interactions between asparagine and arginine/lysine and substrate DNA in the signature [N-(X)_7_-K/R] sequence are conserved. Additionally, interactions from basic amino acids towards substrate binding are also prominent. Perhaps the most notable finding is the involvement of the histidine in DNA binding activity in all cases. As already mentioned, the absence of histidine at the homologous position of plant/opisthokont bZIPs led to further speculation about its possible evolutionary impact on the origin and evolution of all Aureo in the bZIP superfamily.

### 3.5 Exploring the importance of Histidine substitution by *in silico* mutageneis, phylogeny and co-evolution analysis

To further understand the role of histidine at the N-terminal basic region of the Aureo bZIPs, *in-silico* mutagenesis in sequence and structure was conducted. Histidine was mutated to serine (as in plant bZIPs), alanine (as in opisthokont bZIPs) and leucine (as in Aureo2) to find out the consequences in terms of evolution as well as DNA binding respectively. No significant polar interaction or hydrogen bonding is found when His362 is mutated to alanine. When leucine replaces His362/His153 in the docked *Es*Aureo1 / *Sj*Aureo5-DNA complex, no interactions with any of the nucleotides can be seen. His51 in *Es*Aureo3, upon substitution by leucine, remains in close contact with dG’6 of the substrate DNA. Similar situation arises in *Es*Aureo4, where His50 upon mutation to leucine remains in close contact with dG’6. However, no polar contacts could be observed in both these cases. His362 of *Es*Aureo1 on being mutated to serine, comes in close contact with dC’7. While *Es*Aureo2’s Leu197 to serine mutation maintains dT’8 contact, His51→Ser in *Es*Aureo3, shows possible interaction with the N7 of DG’6. Incorporation of serine at the homologous histidine positions in *Es*Aureo4 and *Sj*Aureo5, fails to show any significant interaction. We must acknowledge that the experimental 3D structure of the Aureo-DNA complex could be most reliable. However, our modeled docked structures indicate an active role of His in Aureo-box DNA binding activity. Single residue substitution can influence DNA binding affinity, as noticed in one of our earlier collaborative studies – FD1/FD2 belonging to plant bZIP subfamily A, exhibited a 10-fold difference in DNA binding activity (Dutta et al., 2021). To elaborate further on single histidine substitution, not only we performed MD simulations but also conducted *in vitro* experiments with *Es*Aureo1_wild, *Es*Aureo1_HA and *Es*Aureo1_HS constructs – we discuss the results in the subsequent section of this manuscript. We also wanted to check the evolutionary significance of histidine at this specific position. Therefore, we re-constructed the phylogenetic tree after altering the histidine/leucine position in Aureo by serine (similar to plant bZIPs) and alanine (similar to opisthokont bZIPs). Very interestingly, upon serine substitution, Aureos shift closer to the plant bZIPs sharing a common evolutionary lineage with all the plant bZIP subgroups except D, F and H. All the opisthokont bZIPs are clustered in a different lineage in this case [**Figure-5.a**]. Perhaps this is important to mention here that plant bZIPs D, F and H differ from the remaining plant bZIPs at the crucial 4th/8th position of basic region signature sequence. D and F plant bZIPs possess alanine instead of serine [N(X)_2_<u>A</u>(X)_5_], similar to opisthokont bZIPs [**SI-3**]. F and H bZIPs possess Y and A respectively at the 8th position instead of typical serine [N(X)_6_<u>Y/A</u>R] [**SI-3**]. Upon mutation to alanine, the bZIPs of Aureo aligned in a lineage with opisthokont and plant bZIPs, albeit differently from the wild types. [**Figure-5.b**]. All through we considered a bootstrap value of 1000 to obtain statistically confident data. And indeed, the presence of histidine in Aureo does make a difference in their evolutionary origin. Co-evolution studies conducted with *Es*Aureo1 bZIP with an E-value cut-off 10^-10^ followed by Weblogo analysis shows distinct presence of histidine [**Figure-1.c**] in Aureo subtypes. Upon increasing the E-value to 10^-4^ to accommodate more bZIP sequences however led to complete disappearance of histidine and appearance of serine/alanine at the homologous position [**Figure-5.c**].

**Figure-5:**
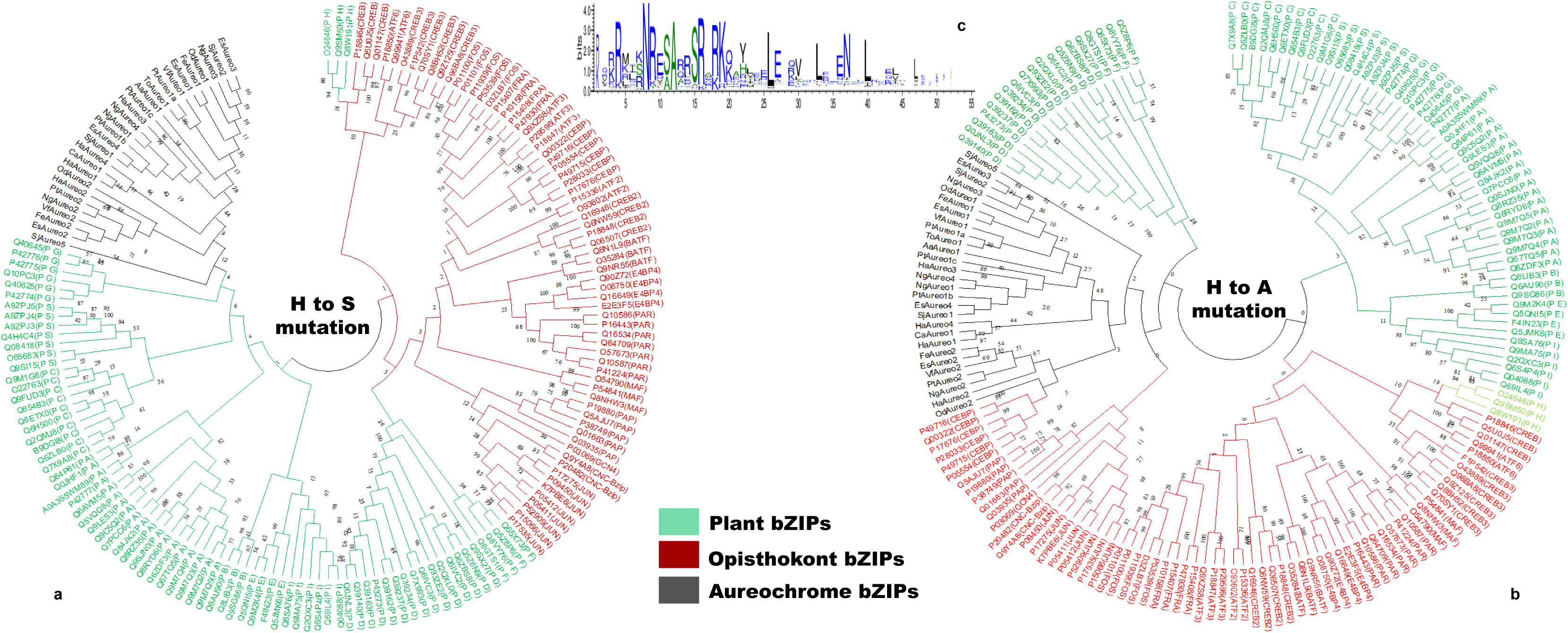
a. Re-construction of phylogenetic tree upon mutation of Aureos’ histidine to serine (plant like) b. Re-construction of phylogenetic tree upon mutation of Aureos’ histidine to alanine (opisthokont like) c. Weblogo result obtained from co-evolution study at E-value cut-off 10^-4.^

### 3.6 Probing the Dynamics of bZIP-DNA Interactions through Molecular Dynamics (MD) Simulations

MD simulations were performed to understand the consistency of interaction between *Es*Aureo1_wild, *Es*Aureo1_HA and *Es*Aureo1_HS and the substrate DNA, albeit over a short time scale. We particularly monitored the behaviour of residue 362 (His, Ala and Ser respectively) throughout the MD run. Thus, the post-simulation analyses conducted include RMSD (<u>R</u>oot <u>M</u>ean <u>S</u>quare <u>D</u>eviation), RMSF (<u>R</u>oot <u>M</u>ean <u>S</u>quare <u>F</u>luctuation), the radius of gyration (Rg), Number of Contacts (within 3 nm cutoff) and the number of hydrogen bonds.

#### 3.6.1. RMSD

The RMSD of the protein backbone (Cɑ-atoms) was analyzed from the simulation data and plotted to compare structural stability across variants. Notably, *Es*Aureo1_wild exhibited consistently lower deviation compared to the mutant variants (*Es*Aureo1_HA and *Es*Aureo1_HS), with average RMSD values of 0.346 nm, 0.395 nm, and 0.414 nm, respectively [**Figure-6.a**]. While all variants showed similar deviation trends at the outset, the wild type demonstrated a marked decrease in deviation around 5 ns, contrasting with the rising trends observed in the mutants. A transient surge in deviation for *Es*Aureo1_wild occurred near 17 ns, followed by alignment with mutant trends between 25 and 30 ns. After a minor fluctuation around 30 ns, the wild type resumed its low-deviation profile, maintaining this trend until the end of the simulation (60 ns). Notably, the *Es*Aureo1_HS variant exhibited a significant increase in RMSD between 50 and 60 ns, stabilizing at a higher deviation. These observations underscore the structural resilience of *Es*Aureo1_wild compared to its mutants.

**Figure 6:**
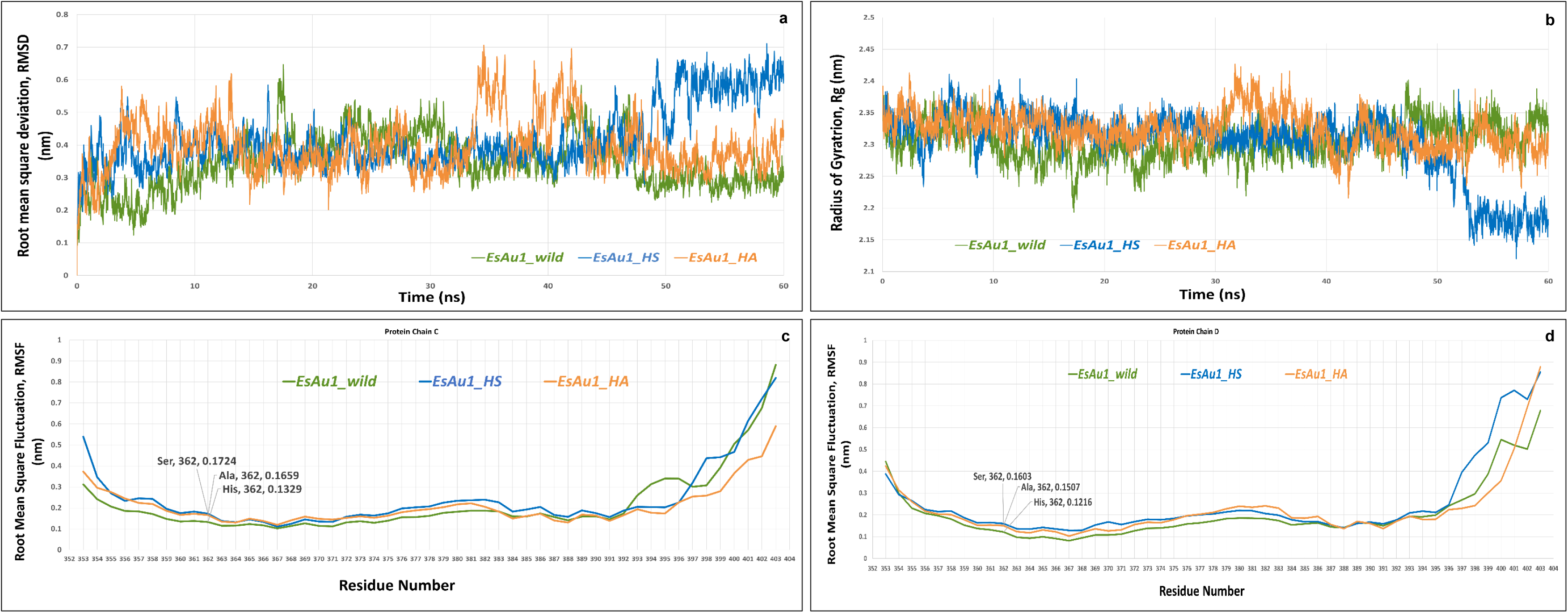
a. Root Mean Square Deviation (RMSD) in nm plotted for *Es*Aureo1_wild (green), *Es*Aureo1_HS (blue) and *Es*Aureo1_HA (orange) protein backbone for 60 ns timescale of MD simulation. b. Radius of Gyration (Rg) in nm for *Es*Aureo1_wild (green), *Es*Aureo1_HS (blue) and *Es*Aureo1_HA (green) plotted against the simulation time of 60 ns. c. Root Mean Square Fluctuation (RMSF) of the protein residues for *Es*Aureo1_wild (orange), *Es*Aureo1_HS (green) and *Es*Aureo1_HA (yellow), as obtained from 60ns simulation data, for Protein chain C. d. Root Mean Square Fluctuation (RMSF) of the protein residues for *Es*Aureo1_wild (orange), *Es*Aureo1_HS (green) and *Es*Aureo1_HA (yellow) for Protein Chain D as obtained from 60 ns simulation data. *[EsAu1 has been used as the abbreviation of EsAureo1]

#### 3.6.2 RMSF

To determine the flexibility of the protein residues, especially His362, the RMSF data was elucidated and plotted against the simulation runtime. A subtle variation, commonly suspected to arise from differences in binding dynamics and solvent interactions, is generally expected between protein chains C and D of the homodimer; hence, they were analyzed and plotted separately [**Figure-6.c,d**]. In both polypeptide chains of Aureo-dimer, His362 of *Es*Aureo1_wild shows the least flexibility, followed by Ala362 of *Es*Aureo1_HA and then Ser362 of *Es*Aureo1_HS.

#### 3.6.3 Radius of gyration

The Rg represents the overall compactness of the protein by measuring the average distance of its atoms from an imaginary axis passing through its center of gravity. Analysis of the Rg curves [**Figure-6.b**] reveals that *Es*Aureo1_wild exhibits greater compactness throughout the simulation compared to its mutant counterparts. Up to the 30 ns mark, all variants display a similar trend, but beyond this point, the wild-type variant maintains a consistency of its Rg, indicating a more compact structure with minimal fluctuation. In contrast, the *Es*Aureo1_HA and *Es*Aureo1_HS mutants show increased fluctuations and less compact conformations over time. Notably, the *Es*Aureo1_HS variant exhibits a sudden and significant drop in Rg after the 50 ns mark, reflecting a collapse in compactness that correlates with the sharp rise in RMSD observed in the same time frame. This behavior underscores the distinct dynamic stability of the wild variant compared to its mutants till the end of the simulation (60 ns).

#### 3.6.4. State of hydrogen bonds

The variation in the number of hydrogen bonds made by His362 with the DNA in both the wild and mutant variants was obtained and was plotted against the simulation time (60000ps). According to the graphs, the residue His362 of *Es*Aureo1_wild shows the maximum number of hydrogen bonds, fluctuating between 0 to 4 [**Figure-7.a**]. The residue Ser362 of *Es*Aureo1_HS shows a moderate number of H-bonds throughout the simulation, fluctuating between 0 to 3 [**Figure-7.b**]. The residue Ala362 of EsAureo1_HA shows no H-bonds throughout the simulation period with the DNA [**Figure-7.c**]. Hydrogen bonding interactions were visualized from the post-MD trajectory files, using PyMOL, with a 10ns interval between each state analyzed for an overall period of 60ns [**Figure-8**].

**Figure 7:**
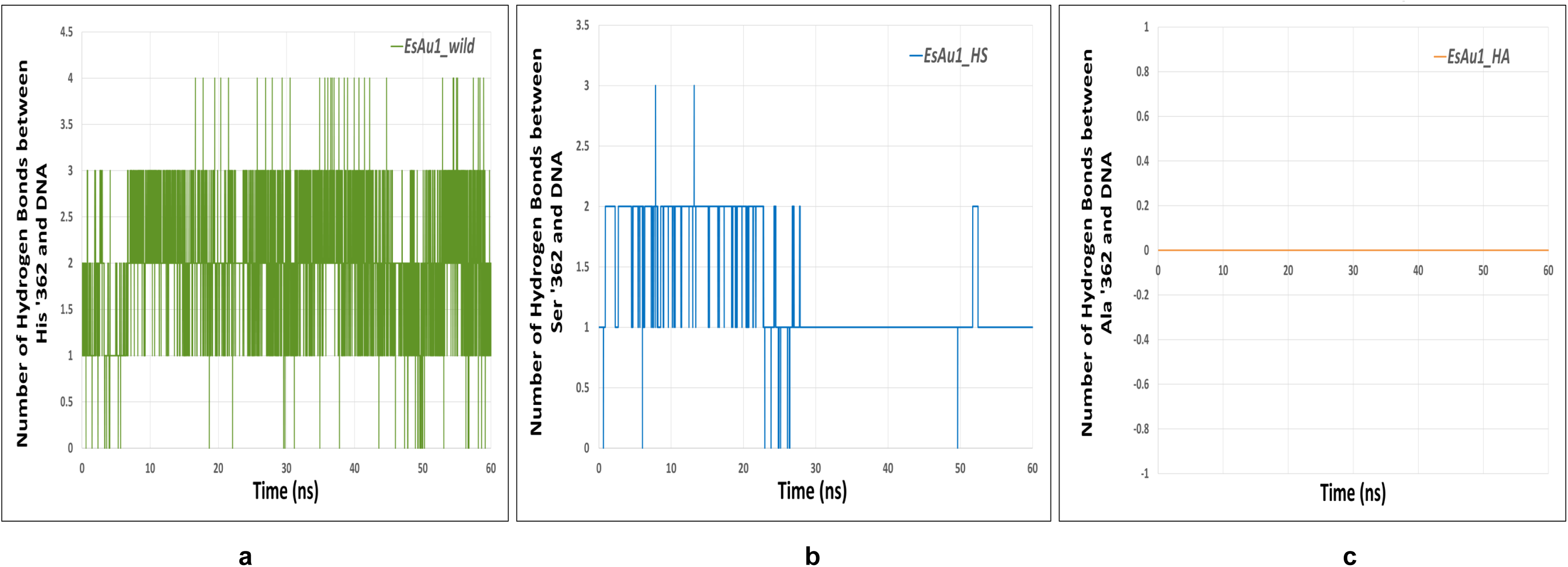
a. Average number of intermolecular hydrogen bonds between His362 in *Es*Aureo1_wild and DNA during 60 ns timescale. b. Average number of intermolecular hydrogen bonds between Ser362 in *Es*Aureo1_HS and DNA c. Average number of intermolecular hydrogen bonds between Ala362 in *Es*Aureo1_HA and DNA. *[EsAu1 has been used as the abbreviation of EsAureo1]

**Figure 8:**
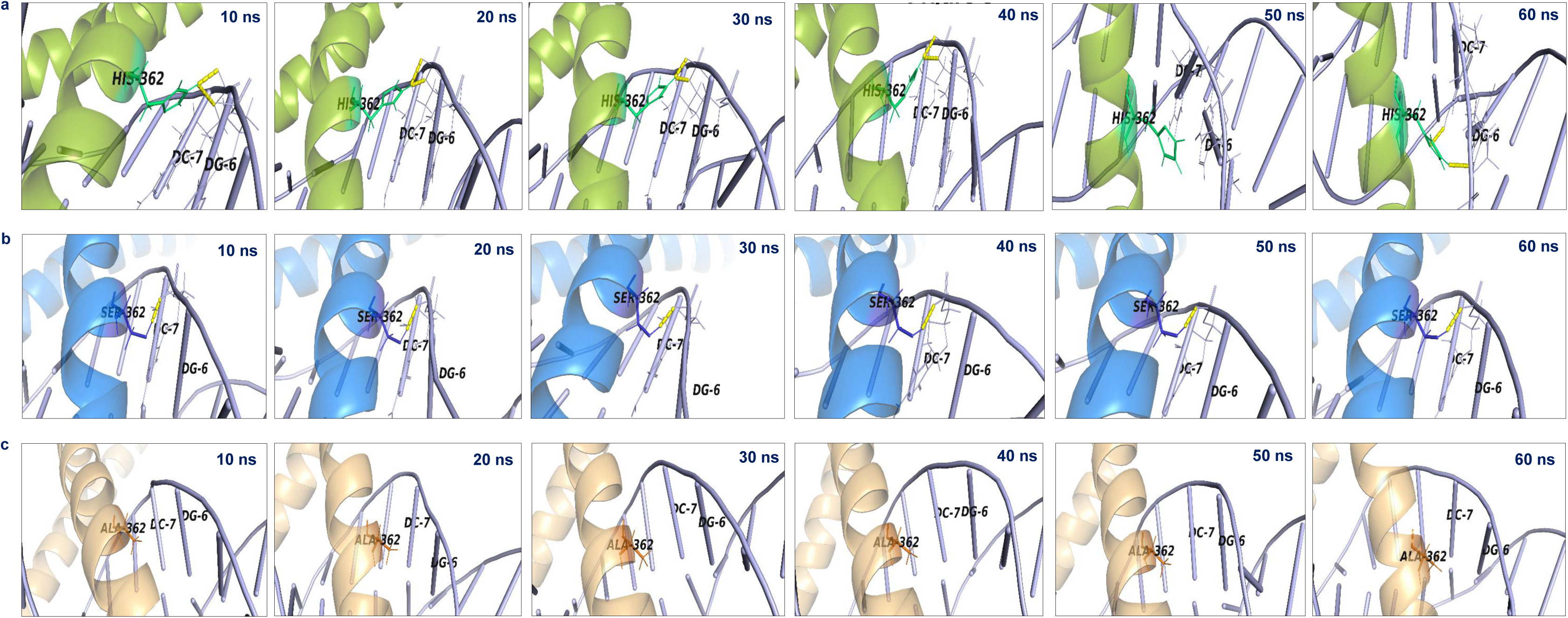
Post-simulation visualization of Residue 362 and its hydrogen bonding interaction with DNA (shown in yellow dashed line), at timesteps of 10, 20, 30, 40, 50 and 60ns (from left to right). a. *Es*Aureo1_wild in green, b. *Es*Aureo1_HS in blue, c. *Es*Aureo1_HA in orange. *[*Es*Au1 has been used as the abbreviation of *Es*Aureo1]

### 3.7. In vitro protein-DNA interaction study by Electrophoretic Mobility Shift Assay

EMSA was performed with the purified and desalted samples of *Es*Aureo1_wild along with its two mutants - *Es*Aureo_HS and *Es*Aureo_HA to compare their DNA binding ability under BL. In all cases, the shift from unbound to substrate DNA bound state was observed with the increase in concentrations of protein [**Figure-9**]. Presence of free DNA bands are indicative of protein being unbound with its substrate DNA. Gradual disappearance of free DNA bands and the resultant shift in electrophoretic mobility is indicative of protein-DNA complex formation. The highest concentration of *Es*Aureo1_wild at which an unbound DNA band was evident first was 0.04 µM [**Figure-9.a**]. In the case of *Es*Aureo_HS [**Figure-9.b**] and *Es*Aureo_HA [**Figure-9.c**], the first presence of an unbound DNA band showed up at concentrations 0.076 µM and 0.082 µM respectively. Hence, although all of the variants manifest DNA binding, the result of EMSA indicates that *Es*Aureo1_wild has a higher DNA binding affinity. And the unique presence of histidine at the basic region of Aureo bZIPs seems justifiable for their specific interaction with Aureo-box substrates.

**Figure 9:**
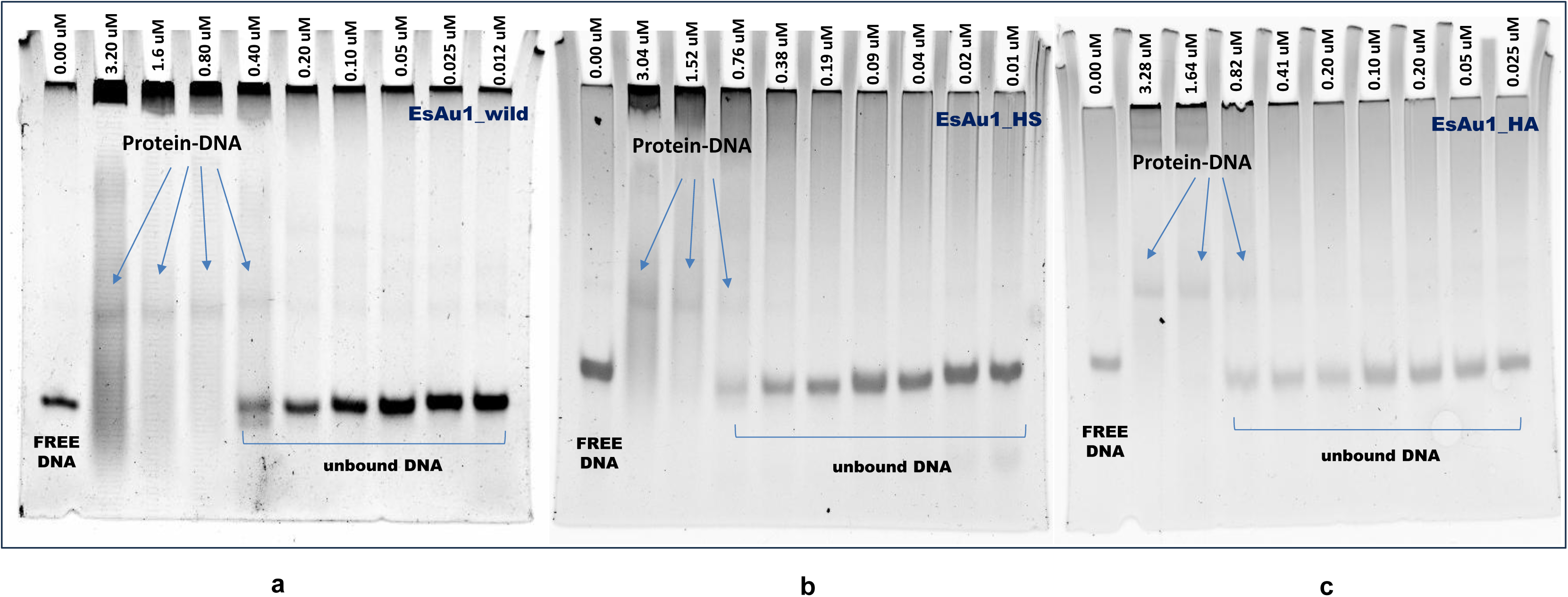
*In vitro* protein□DNA interaction study of Aureochrome_bZIPs through electrophoretic mobility shift assay-a. *Es*Aureo1_wild, b. *Es*Aureo_HS, c. *Es*Aureo_HA.

## 4. Discussion

The global network of TFs comprising interactions with specific substrate DNA or partner protein molecules regulates one of the most fundamental processes of life - gene expression. Besides the possibility of simple/auto-regulation, a TF can participate in feed forward loops/feedback loops (FFLs/FBLs)(Alon, 2007) as homodimer or heterodimer, in complex and dynamic gene expression regulatory units [**Figure-10**]. While the orderliness and sequence composition of the basic and fork regions of bZIPs play a crucial role in DNA binding specificity, sequence organization and stability of heptad repeats in the zipper region is equally important. Even a subtle change in heptad sequence can impact dimerization specificity and stability, and the resulting manifestation could be significant. Choosing partners from the same or different group of bZIPs, respectively for homo- and hetero-dimerization can generate substantial functional diversity. The 3D structure of a protein can be considered as a network of nodes (amino acid residues) connected via edges (non-covalent interactions) and the relative importance of the individual amino acid residues can be obtained by calculating different topological parameters of RIG. Therefore, apart from characterizing protein-protein/DNA interface in terms of bonding interactions, surface area and energy, we conducted all-atom network analysis. Centrality measures like degree, closeness and betweenness effectively captures influential nodes in a network. In this study, we emphasized on measuring EC values considering its potential to detect nodes important for making connections and information flow, especially allostery. Interestingly, residues with highest EC values are always at ‘a’ and ‘d’ positions of a heptad in all eight representative co-crystal structures. Importance of hydrophobic residues at the strategic positions necessary for dimerization and consequent enhancement of DNA binding activity has been shown for c-Fos mutants(Porte et al., 1997). Incorporation of isoleucine at the 2nd heptad not only promoted homodimerization but also conferred DNA binding specificity towards TRE substrate in c-Fos mutants. While EC calculations for Aureos await determination of experimental 3D structures, sequence comparison does indicate that Aureo bZIPs are mostly similar to other bZIPs at least in these specific nodes except ‘d’ residue in heptad 4. Measurement of EC undoubtedly reiterates the importance of hydrophobic residues towards maintaining stable bZIP dimerization, signal transfer and thereby promoting DNA binding activity. Therefore, insights obtained from EC measurements could be helpful for designing synthetic sensory TF in the near future.

**Figure 10:**
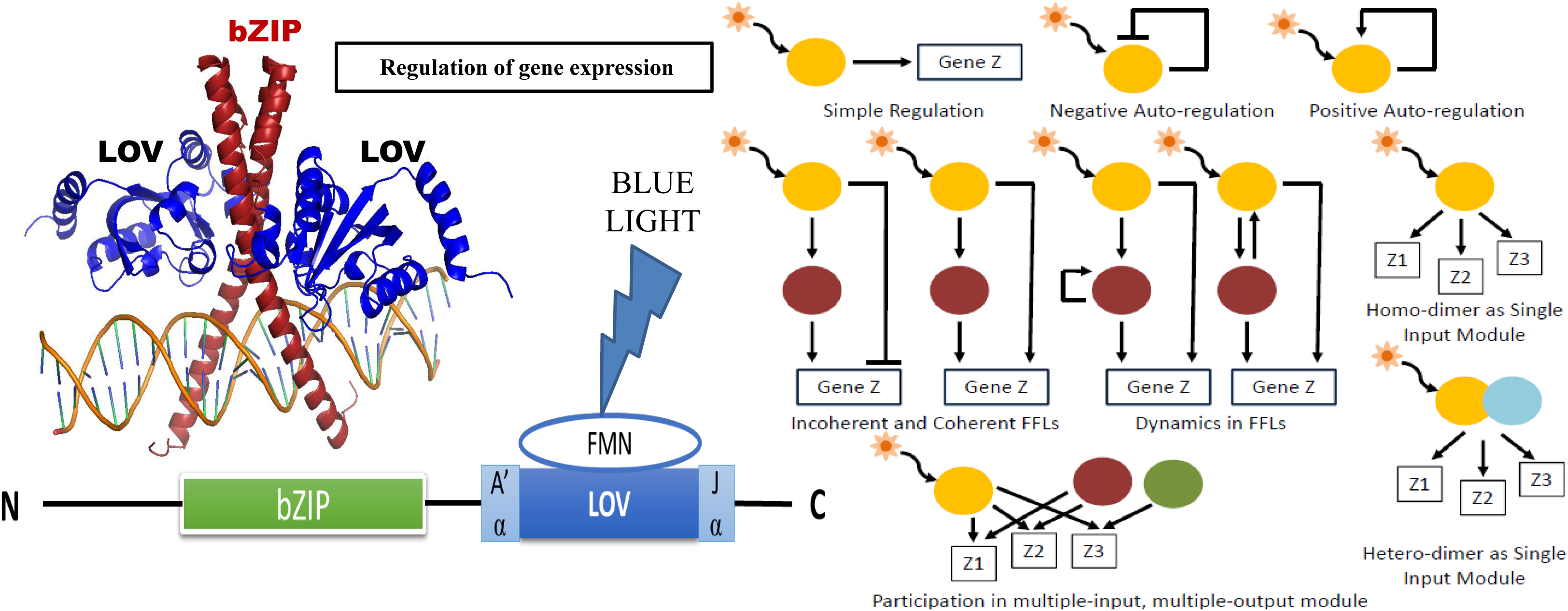
Cartoon and diagrammatic representation of the sensory transcription regulation network, where Aureochrome is perceived to act as a sensory, blue-light regulated transcription factor regulating gene expression. The concept of sensory transcription regulation network is adopted from Network motifs: theory and experimental approaches by Alon U. [ref. 56, doi:10.1038/nrg2102]

The BL-responsive sensory TFs like Aureo, the pivot of this study, comes with an additional advantage. Such an association of one major group of eukaryotic TF (here, bZIPs) with a light sensor (here, LOV) is rare and fascinating. Our study therefore attempts to draw a comparative analysis of Aureos and the other conventional bZIP proteins without LOV sensors. With distinct differences noticed, it remains to be clarified whether or not these variations arose through evolutionary acquisition of the sensor domain to confer physiological advantages to the host organisms. Though bZIPs are supposed to have far reaching origin, the expansion and diversification of the bZIP superfamily has been found to be positively correlated with the evolution of eukaryotes (Jindrich & Degnan, 2016). The number of bZIP homologs as well as the complexity of bZIPs have increased over the time in the process of evolution. Initially bZIPs were supposed to evolve independently in 4 eukaryotic lineages – planta, opisthokonta, amoebozoa, and heterokonta (Jindrich & Degnan, 2016). Among them 17 bZIP subfamilies within opisthokont, namely, YAP, MAF, XBP1, CNC, BATF, CREB, CREB-2, OASIS, PAR, E4BP4, ATF-2, ATF-4, ATF-6, FOS, JUN, C/EBP, PAP (Amoutzias et al., 2007) (Fujii et al., 2000) and ten distinct subclasses (A, C, D, G and S) of plant bZIPs (Jakoby et al., 2002) (Corrêa et al., 2008) are evident. Whereas in heterokontophyta (**SI-7]**, the phylum comprising of 12 families under the algal group-4 as proposed in the algal classification by Robert Lee (Lee, 2018), the most abundant bZIP transcription factor is Aureo (Rayko et al., 2010). It has been observed in the evolutionary analysis performed by us that all the bZIPs of different Aureo homologs together form a different clade albeit shared the ancestor with opishtokont bZIP subfamilies-JUN, MAF, CREB, CREB3, PAP, GCN4 and plant bZIP subfamilies. But substitution of histidine results in an assembly of Aureo-bZIPs predominantly with plant bZIPs sharing a common ancestral lineage. This study therefore investigates how nuances at the residue level are manifested in the phyletic distribution of bZIP members, including the evolution of Aureos and their subtypes.

The bZIPs of the metazoans recognize 6 different consensus DNA-binding sites- <u>c</u>-AMP responsive <u>e</u>lement (CRE- TGACGTCA), <u>T</u>PA responsive <u>e</u>lement (TRE- ATGAGTCAT), CAAT box (TTGCGCAA), PAR binding sites (TTACGTAAT), <u>Ma</u>f recognition <u>e</u>lement (MARE- CTGATCAGC) and CRE-like elements (TGACGTGC) (Amoutzias et al., 2007) (Rodríguez-Martínez et al., 2017). Again, CRE, TRE and YAP (<u>y</u>east <u>a</u>ctivator <u>p</u>rotein) sites can be recognized by fungal bZIPs. Plant bZIPs recognize G-box (CACGTG), A-box (TACGTA), C-box (GACGTC), S-box (TGACGTG), D-box (TGACGTG) (Wang et al., 2018) etc. We conducted *in-silico* and *in-vitro* studies, using Aureo-box containing CRE (TGACGTCA) as the substrate DNA [sequence taken from X-ray structure of CREB bZIP- CRE complex, PDB ID: 1DH3]. The docking, MD simulation as well as EMSA studies were performed not only to check the DNA binding efficiency of the Aureo_bZIPs (wild type and mutants) but also to explore the precise role of histidine in DNA binding. Both theory and experiments indicate significance of histidine at the basic region of aureochrome. It can be concluded that His362 plays a key role in positioning the *Es*Aureo1 bZIP to the core ACGT sequence by favorably interacting with its flanking sequence. Changing this amino acid to Ser (as in plants) and Ala (as in opisthokonts) leads to a significant decrease in polar interactions [**Figure-7,8**] [**SI-8**] and therefore impacts the stability of the complex [**Figure-6.a**] [as found from MD Simulation study]. However, at the same time, the overall binding strength of all the constructs remains comparable [as confirmed by the EMSA studies] [**Figure-9**]. This makes it clearer that His is very strongly related to the Aureo box, and thus the slightest change in this recognition box (as seen in opisthokonts and plants) makes this amino acid residue site prone to change. This provides an answer for the causality of co-evolution. It is worth reiterating that Aureos are light-responsive TFs and therefore should function like a photo-switch. Studies regarding the kinetics of binding and the width of spectrum of the changes in functionality of the bZIP due to variation in substrate DNA sequence, are still an area requiring further work. The binding specificity of a TF to DNA is intricately governed by its interaction with the minor groove geometry and regional electrostatics. Among the three mutants analyzed through simulation, *Es*Aureo1_wild has the maximum DNA binding ability with the highest number of hydrogen bonds present at the bZIP-DNA interface. *Es*Aureo1_HS ranks in between wild type and *Es*Aureo1_HA. The alanine mutant modeled after opisthokont bZIP proteins, emerged as distinct, with Ala362 failing to interact with the flanking residues of the ACGT core. While the number of non-covalent interactions, e.g. hydrogen bonds, determines DNA binding specificity(Lin & Guo, 2019), prompt and selective binding is eventually reflected in protein-DNA association kinetics. Molecular docking, MD simulation as well as EMSA studies reveal that the placement of histidine bespoke ‘Aureo-box’ specific interaction. However, it would be highly interesting to see whether histidine makes any difference in association kinetics of Aureos towards substrate recognition/binding upon BL illumination. Strength and specificity of DNA binding interactions as well as variable association kinetics can be adopted further for developing precision optogenetic tools.

Perhaps another important observation that we made in this study is the unique presence of histidine in all Aureo bZIPs, except Aureo2 from several stramenopiles. Moreover, Aureo2 also possesses non-functional LOV (Takahashi et al., 2007). Out of eleven conserved amino acids in the flavin-binding pocket, nine are conserved in Aureo2. The reason behind such differences in Aureo2 is elusive – it is possible that the sensor domain of Aureo2 could be an intermediate between non-light sensing PAS (<u>P</u>er-<u>A</u>RNT-<u>S</u>im, ancestor to LOV) and LOV. Our study further unveiled that histidine co-evolves with serine at the 8th position of the signature bZIP sequence. Interestingly in Aureo2, both 4th and 8th positions are differently occupied by leucine and threonine respectively. Despite such marked changes in the basic region, Aureo2 clubs with the rest Aureos in the phylogenetic tree and not with plants/opisthokonts. However, upon single substitution from leucine to alanine, Aureo2 shifts lineage and sides with CEBPs of opisthokonts, while retaining proximity with other Aureos. Considering the presence of non-functional sensor domain and close association with opisthokont bZIPs just upon single amino acid substitution, we speculate that Aureo2 might have evolved first among the Aureos. During the course of evolution, possibly with the gain of function mutation, Aureos have finally acquired the ability to sense light and perform light dependent transcriptional activities. And perhaps to preserve this very important property, several other Aureo paralogs emerged through gene duplication events.

In summary, the bZIPs of aureochromes share close structural and functional similarities with other bZIP TFs found across plants and animals, exhibiting comparable dimerization properties and DNA-binding affinities to substrate DNA. Consequently, this bZIP integrated with its blue light sensor, can be utilized in plants and animals either as a genetically encoded recombinant protein driven by a synthetic promoter or as a transgene expressed under native promoters in target cells to achieve tissue-specific gene regulation via light stimulation. Furthermore, orthogonal manipulation through engineered gene regulatory circuits can facilitate non-invasive control over existing regulatory networks. Our previous study indicates that the aureochrome-bZIP monomer dimerizes with bZIP monomers within the aureochrome family as well as with other non-aureochrome bZIPs (Khamaru et al., 2024). As such, introducing light-sensitive aureochrome-bZIPs into genetically modified transgenic lines or transiently transformed cells, can enable light-mediated regulation of developmental pathways and stress responses by interacting with the inherent bZIP proteins of plants or animals. Additionally, all the bZIP proteins regulate crucial developmental processes of plants and animals like-growth, cell division, and differentiation can be further modulated by adjusting the duration of gene activation or repression through fine tuning of the light sensor module associated with these bZIPs. Interestingly, all the subclasses of plant bZIP proteins that bind to promoter or regulator present upstream of the target genes, recognize the ACGT sequence of DNA. As Aureo-bZIPs also recognizes ACGT sequences, it seems to be a promising optogenetic tool for regulating plant gene expression and influencing the existing bZIP dependent metabolic and developmental pathways within plant kingdom. Thus, characterization of these exclusive light regulated bZIPs not only gives insight to the transcriptional regulation pattern of the lower group of plants i.e. the heterokont algae, but shed light into the scope of optogenetic regulation of plant genes.

## Acknowledgement

MK and AD acknowledge University Grants Commission (UGC) and Department of Biotechnology (DBT), Government of India respectively, for their doctoral research fellowships. DM acknowledges funding from DBT, Government of India (BT/PR26435/BRB/10/1627/2017) for supporting the research work during the initial phase and SERB, Govt of India (CRG/2023/006463) at the present phase. The authors acknowledge the central instrumentation facility as well as the computer facility of the department supported by DBT-BUILDER and DST-FIST programs respectively. The authors also thank Soumen Roy, Bose Institute - Kolkata for important discussions on network analysis.

## Author contribution

**Madhurima Khamaru**: Data curation, Investigation, Methodology, Formal analysis, Validation, Visualization, Writing – original draft, Writing – review and editing; **Debarshi Bose**: Investigation, Methodology, Formal analysis, Visualization, Writing – original draft; **Anwesha Deb**: Investigation, Methodology; **Devrani Mitra**: Conceptualization, Supervision, Funding acquisition, Project administration, Investigation, Methodology, Formal analysis, Validation, Visualization, Writing – original draft, Writing – review and editing.

## Data availability statement

All data generated or analyzed during this study are included in the main text article as well as in the supplementary information. Further enquiries can be directed to the corresponding author.

## References

Alon, U. (2007). Network motifs: theory and experimental approaches. Nature Reviews Genetics 2007 8:6, 8(6), 450–461. 10.1038/nrg2102

Altschul, S. F., Gish, W., Miller, W., Myers, E. W., & Lipman, D. J. (1990). Basic local alignment search tool. Journal of Molecular Biology, 215(3), 403–410. 10.1016/S0022-2836(05)80360-2

Amoutzias, G. D., Veron, A. S., Weiner, J., Robinson-Rechavi, M., Bornberg-Bauer, E., Oliver, S. G., & Robertson, D. L. (2007). One billion years of bZIP transcription factor evolution: Conservation and change in dimerization and DNA-binding site specificity. Molecular Biology and Evolution, 24(3), 827–835. 10.1093/molbev/msl211

Apweiler, R., Bairoch, A., Wu, C. H., Barker, W. C., Boeckmann, B., Ferro, S., Gasteiger, E., Huang, H., Lopez, R., Magrane, M., Martin, M. J., Natale, D. A., O’Donovan, C., Redaschi, N., & Yeh, L. S. L. (2018). UniProt: the universal protein knowledgebase. Nucleic Acids Research, 46(5), 2699–2699. 10.1093/NAR/GKY092

Bader, A. G., & Vogt, P. K. (2006). Leucine Zipper Transcription Factors: bZIP Proteins. In Encyclopedic Reference of Genomics and Proteomics in Molecular Medicine (pp. 964–967). Springer Berlin Heidelberg. 10.1007/3-540-29623-9_2180

Banerjee, A., Herman, E., Kottke, T., & Essen, L. O. (2016). Structure of a Native-like Aureochrome 1a LOV Domain Dimer from Phaeodactylum tricornutum. Structure, 24(1), 171–178. 10.1016/j.str.2015.10.022

Banerjee, A., Herman, E., Serif, M., Maestre-Reyna, M., Hepp, S., Pokorny, R., Kroth, P. G., Essen, L.-O., & Kottke, T. (2016). Allosteric communication between DNA-binding and light-responsive domains of diatom class I aureochromes. Nucleic Acids Research, 44(12), 5957– 5970. 10.1093/nar/gkw420

Banerjee, S., & Mitra, D. (2020). Structural Basis of Design and Engineering for Advanced Plant Optogenetics. Trends in Plant Science, 25(1), 35–65. 10.1016/j.tplants.2019.10.002

Boral, A., Khamaru, M., & Mitra, D. (2022). Designing synthetic transcription factors: A structural perspective. Advances in Protein Chemistry and Structural Biology, 130. 10.1016/BS.APCSB.2021.12.003

Busch, S. J., & Sassone-Corsi, P. (1990). Dimers, leucine zippers and DNA-binding domains. Trends in Genetics. 10.1016/0168-9525(90)90071-D

Cock, P. J. A., Antao, T., Chang, J. T., Chapman, B. A., Cox, C. J., Dalke, A., Friedberg, I., Hamelryck, T., Kauff, F., Wilczynski, B., & De Hoon, M. J. L. (2009). Biopython: freely available Python tools for computational molecular biology and bioinformatics. Bioinformatics (Oxford, England), 25(11), 1422–1423. 10.1093/BIOINFORMATICS/BTP163

Corrêa, L. G. G., Riaño-Pachón, D. M., Schrago, C. G., Vicentini dos Santos, R., Mueller-Roeber, B., & Vincentz, M. (2008). The Role of bZIP Transcription Factors in Green Plant Evolution: Adaptive Features Emerging from Four Founder Genes. PLoS ONE, 3(8), e2944. 10.1371/journal.pone.0002944

Costa, B., Sachse, M., Jungandreas, A., Bartulos, C. R., & Gruber, A. (2013). Aureochrome 1a Is Involved in the Photoacclimation of the Diatom Phaeodactylum tricornutum. PLoS ONE, 8(9). 10.1371/journal.pone.0074451

Cruz, P., Paredes, N., Asela, I., Kolimi, N., Molina, J. A., Ramírez-Sarmiento, C. A., Goutam, R., Huang, G., Medina, E., & Sanabria, H. (2023). Domain tethering impacts dimerization and DNA-mediated allostery in the human transcription factor FoxP1. The Journal of Chemical Physics, 158(19). 10.1063/5.0138782

Deb, A., Grewal, R. K., Roy, S., & Mitra, D. (2020). Residue interaction dynamics in Vaucheria aureochrome1 light-oxygen-voltage: Bridging theory and experiments. Proteins, 88(12), 1660– 1674. 10.1002/PROT.25984

Delano, W. (2002). The PyMOL Molecular Graphics System.

Deppmann, C. D., Acharya, A., Rishi, V., Wobbes, B., Smeekens, S., Taparowsky, E. J., & Vinson, C. (2004). Dimerization specificity of all 67 B-ZIP motifs in Arabidopsis thaliana : a comparison to Homo sapiens B-ZIP motifs. Nucleic Acids Research, 32(11), 3435–3445. 10.1093/NAR/GKH653

Duan, L., Hope, J. M., Guo, S., Ong, Q., François, A., Kaplan, L., Scherrer, G., & Cui, B. (2018). Optical Activation of TrkA Signaling. ACS Synthetic Biology, 7(7), 1685–1693. 10.1021/ACSSYNBIO.8B00126/SUPPL_FILE/SB8B00126_SI_001.PDF

Dutta, S., Deb, A., Biswas, P., Chakraborty, S., Guha, S., Mitra, D., Geist, B., Schäffner, A. R., & Das, M. (2021). Identification and functional characterization of two bamboo FD gene homologs having contrasting effects on shoot growth and flowering. Scientific Reports, 11(1). 10.1038/S41598-021-87491-6

Evgeny, K. (2010). Crystal contacts as nature’s docking solutions. Journal of Computational Chemistry, 31(1), 133–143. 10.1002/JCC.21303

Fujii, Y., Shimizu, T., Toda, T., Yanagida, M., & Hakoshima, T. (2000). letters Structural basis for the diversity of DNA recognition by bZIP transcription factors.

Glover, J. N. M., & Harrison, S. C. (1995). Crystal structure of the heterodimeric bZIP transcription factor c-Fos–c-Jun bound to DNA. In Nature. 10.1038/373257a0

Grigoryan, G., & Keating, A. E. (2006). Structure-based Prediction of bZIP Partnering Specificity. Journal of Molecular Biology, 355(5), 1125–1142. 10.1016/j.jmb.2005.11.036

Hagberg, A. A., Schult, D. A., & Swart, P. J. (2008). Exploring network structure, dynamics, and function using NetworkX. 7th Python in Science Conference (SciPy 2008), 11–15. https://www.bibsonomy.org/bibtex/272096c6553d7057409fe78ca698eb332/mschuber

Hakoshima, T. (2005a). Leucine Zippers. In Encyclopedia of Life Sciences. John Wiley & Sons, Ltd. 10.1038/npg.els.0005049

Hakoshima, T. (2005b). Leucine Zippers. In Encyclopedia of Life Sciences. 10.1038/npg.els.0005049

Harbury, P. B., Zhang, T., Kim, P. S., & Alber, T. (1993). A switch between two-, three-, and four-stranded coiled coils in GCN4 leucine zipper mutants. Science (New York, N.Y.), 262(5138), 1401–1407. 10.1126/SCIENCE.8248779

Heintz, U., & Schlichting, I. (2016). Blue light-induced LOV domain dimerization enhances the affinity of aureochrome 1a for its target DNA sequence. ELife, 5(JANUARY2016). 10.7554/ELIFE.11860

Hepp, S., Trauth, J., Hasenj, S., Bezold, F., Essen, L.-O., & Taxis, C. (2020). An Optogenetic Tool for Induced Protein Stabilization Based on the Phaeodactylum tricornutum Aureochrome 1a LighteOxygeneVoltage Domain. 10.1016/j.jmb.2020.02.019

Hepp, S., Trauth, J., Hasenjäger, S., Bezold, F., Essen, L. O., & Taxis, C. (2020). An Optogenetic Tool for Induced Protein Stabilization Based on the Phaeodactylum tricornutum Aureochrome 1a Light–Oxygen–Voltage Domain. Journal of Molecular Biology. 10.1016/j.jmb.2020.02.019

Huang, Y., Wang, L., Zheng, M., Zheng, L., Tong, Y., & Li, Y. (2014). Overexpression of NgAUREO1, the gene coding for aurechrome 1 from Nannochloropsis gaditana, into Saccharomyces cerevisiae leads to a 1.6-fold increase in lipid accumulation. Biotechnology Letters, 36(3), 575–579. 10.1007/s10529-013-1386-0

Huysman, M. J. J., Fortunato, A. E., Matthijs, M., Costa, B. S., Vanderhaeghen, R., Van den Daele, H., Sachse, M., Inzé, D., Bowler, C., Kroth, P. G., Wilhelm, C., Falciatore, A., Vyverman, W., & De Veylder, L. (2013). AUREOCHROME1a-Mediated Induction of the Diatom-Specific Cyclin dsCYC2 Controls the Onset of Cell Division in Diatoms (Phaeodactylum tricornutum). The Plant Cell, 25(1), 215–228. 10.1105/tpc.112.106377

Jakoby, M., Weisshaar, B., Dröge-Laser, W., Vicente-Carbajosa, J., Tiedemann, J., Kroj, T., & Parcy, F. (2002). bZIP transcription factors in Arabidopsis. Trends in Plant Science, 7(3), 106–111. 10.1016/S1360-1385(01)02223-3

Jerng, H. H., Patel, J. M., Khan, T. A., Arenkiel, B. R., & Pfaffinger, P. J. (2021). Light-regulated voltage-gated potassium channels for acute interrogation of channel function in neurons and behavior. PLOS ONE, 16(3), e0248688. 10.1371/JOURNAL.PONE.0248688

Ji, C., Mao, X., Hao, J., Wang, X., Xue, J., Cui, H., & Li, R. (2018). Analysis of bZIP transcription factor family and their expressions under salt stress in Chlamydomonas reinhardtii. International Journal of Molecular Sciences, 19(9). 10.3390/ijms19092800

Jindrich, K., & Degnan, B. M. (2016). The diversification of the basic leucine zipper family in eukaryotes correlates with the evolution of multicellularity Genome evolution and evolutionary systems biology. BMC Evolutionary Biology, 16(1), 1–12. 10.1186/s12862-016-0598-z

Kamisetty, H., Ovchinnikov, S., & Baker, D. (2013). Assessing the utility of coevolution-based residue-residue contact predictions in a sequence- and structure-rich era. Proceedings of the National Academy of Sciences of the United States of America, 110(39), 15674–15679. 10.1073/PNAS.1314045110/SUPPL_FILE/SD05.XLSX

Khamaru, M., Nath, D., Mitra, D., & Roy, S. (2024). Assessing Combinatorial Diversity of Aureochrome Basic Leucine Zippers through Genome-Wide Screening. Cells, Tissues, Organs, 213(2), 133–146. 10.1159/000527593

Kroth, P. G., Wilhelm, C., & Kottke, T. (2017). An update on aureochromes_ Phylogeny – mechanism – function. Journal of Plant Physiology, 217, 20–26. 10.1016/j.jplph.2017.06.010

Krylov, D., Mikhailenko, I., & Vinson, C. (1994). A thermodynamic scale for leucine zipper stability and dimerization specificity: e and g interhelical interactions. The EMBO Journal, 13(12), 2849. 10.1002/J.1460-2075.1994.TB06579.X

Kumar, S., Stecher, G., Li, M., Knyaz, C., & Tamura, K. (2018). MEGA X: Molecular Evolutionary Genetics Analysis across Computing Platforms. Molecular Biology and Evolution, 35(6), 1547– 1549. 10.1093/MOLBEV/MSY096

Lee, R. E. (2018). Phycology. Phycology. 10.1017/9781316407219

Lin, M., & Guo, J. T. (2019). New insights into protein–DNA binding specificity from hydrogen bond based comparative study. Nucleic Acids Research, 47(21), 11103–11113. 10.1093/NAR/GKZ963

Mann, M., Serif, M., Wrobel, T., Eisenhut, M., Madhuri, S., Flachbart, S., Weber, A. P. M., Lepetit, B., Wilhelm, C., & Kroth, P. G. (2020). The Aureochrome Photoreceptor PtAUREO1a Is a Highly Effective Blue Light Switch in Diatoms. IScience, 23(11), 101730. 10.1016/J.ISCI.2020.101730

Matiiv, A. B., & Chekunova, E. M. (2018). Aureochromes – Blue Light Receptors. Biochemistry (Moscow), 83(6), 662–673. 10.1134/S0006297918060044

Mitra, D., Yang, X., & Moffat, K. (2012). Crystal structures of aureochrome1 LOV suggest new design strategies for optogenetics. Structure. 10.1016/j.str.2012.02.016

Negre, C. F. A., Morzan, U. N., Hendrickson, H. P., Pal, R., Lisi, G. P., Patrick Loria, J., Rivalta, I., Ho, J., & Batista, V. S. (2018). Eigenvector centrality for characterization of protein allosteric pathways. Proceedings of the National Academy of Sciences of the United States of America, 115(52), E12201–E12208. 10.1073/PNAS.1810452115

Pettersen, E. F., Goddard, T. D., Huang, C. C., Couch, G. S., Greenblatt, D. M., Meng, E. C., & Ferrin, T. E. (2004). UCSF Chimera—A visualization system for exploratory research and analysis. Journal of Computational Chemistry, 25(13), 1605–1612. 10.1002/JCC.20084

Podust, L. M., Krezel, A. M., & Kim, Y. (2001). Crystal structure of the CCAAT box/enhancer-binding protein beta activating transcription factor-4 basic leucine zipper heterodimer in the absence of DNA. The Journal of Biological Chemistry, 276(1), 505–513. 10.1074/JBC.M005594200

Pogenberg, V., Consani Textor, L., Vanhille, L., Holton, S. J., Sieweke, M. H., & Wilmanns, M. (2014). Design of a bZip Transcription Factor with Homo/Heterodimer-Induced DNA-Binding Preference. Structure, 22(3), 466–477. 10.1016/j.str.2013.12.017

Porte, D., Oertel-Buchheit, P., John, M., Granger-Schnarr, M., & Schnarr, M. (1997). DNA binding and transactivation properties of Fos variants with homodimerization capacity. Nucleic Acids Research, 25(15), 3026–3033. 10.1093/NAR/25.15.3026

Pronk, S., Páll, S., Schulz, R., Larsson, P., Bjelkmar, P., Apostolov, R., Shirts, M. R., Smith, J. C., Kasson, P. M., Van Der Spoel, D., Hess, B., & Lindahl, E. (2013). GROMACS 4.5: a high-throughput and highly parallel open source molecular simulation toolkit. Bioinformatics, 29(7), 845–854. 10.1093/BIOINFORMATICS/BTT055

Rayko, E., Maumus, F., Maheswari, U., Jabbari, K., & Bowler, C. (2010). Transcription factor families inferred from genome sequences of photosynthetic stramenopiles. The New Phytologist, 188(1), 52–66. 10.1111/J.1469-8137.2010.03371.X

Rodríguez-Martínez, J. A., Reinke, A. W., Bhimsaria, D., Keating, A. E., & Ansari, A. Z. (2017). Combinatorial bZIP dimers display complex DNA-binding specificity landscapes. ELife, 6. 10.7554/eLife.19272

Schumacher, M. A., Goodman, R. H., & Brennan, R. G. (2000). The Structure of a CREB bZIPSomatostatin CRE Complex Reveals the Basis for Selective Dimerization and Divalent Cation-enhanced DNA Binding*. 10.1074/jbc.M007293200

Sievers, F., Wilm, A., Dineen, D., Gibson, T. J., Karplus, K., Li, W., Lopez, R., McWilliam, H., Remmert, M., Söding, J., Thompson, J. D., & Higgins, D. G. (2011). Fast, scalable generation of high-quality protein multiple sequence alignments using Clustal Omega. Molecular Systems Biology, 7(1), 539. 10.1038/MSB.2011.75

Sinden, R. R. (1994). DNA–Protein Interactions. DNA Structure and Function, 287–325. 10.1016/B978-0-08-057173-7.50013-4

Takahashi, F., Yamagata, D., Ishikawa, M., Fukamatsu, Y., Ogura, Y., Kasahara, M., Kiyosue, T., Kikuyama, M., Wada, M., & Kataoka, H. (2007). AUREOCHROME, a photoreceptor required for photomorphogenesis in stramenopiles. Proceedings of the National Academy of Sciences, 104(49), 19625–19630. 10.1073/pnas.0707692104

Tuszynska, I., Magnus, M., Jonak, K., Dawson, W., & Bujnicki, J. M. (2015). NPDock: a web server for protein–nucleic acid docking. Nucleic Acids Research, 43(Web Server issue), W425. 10.1093/NAR/GKV493

Van Der Spoel, D., Lindahl, E., Hess, B., Groenhof, G., Mark, A. E., & Berendsen, H. J. C. (2005). GROMACS: Fast, flexible, and free. Journal of Computational Chemistry, 26(16), 1701–1718. 10.1002/JCC.20291

Vinson, C., Myakishev, M., Acharya, A., Mir, A. A., Moll, J. R., & Bonovich, M. (2002). Classification of Human B-ZIP Proteins Based on Dimerization Properties. Molecular and Cellular Biology, 22(18), 6321–6335. 10.1128/MCB.22.18.6321-6335.2002

Vinson, C. R., Hai, T., & Boyd, S. M. (1993). Dimerization specificity of the leucine zipper-containing bZIP motif on DNA binding: Prediction and rational design. Genes and Development, 7(6), 1047–1058. 10.1101/gad.7.6.1047

Wang, Y., Zhang, Y., Zhou, R., Dossa, K., Yu, J., Li, D., Liu, A., Mmadi, M. A., Zhang, X., & You, J. (2018). Identification and characterization of the bZIP transcription factor family and its expression in response to abiotic stresses in sesame. PLOS ONE, 13(7), e0200850. 10.1371/journal.pone.0200850

Waterhouse, A., Bertoni, M., Bienert, S., Studer, G., Tauriello, G., Gumienny, R., Heer, F. T., De Beer, T. A. P., Rempfer, C., Bordoli, L., Lepore, R., & Schwede, T. (2018). SWISS-MODEL: homology modelling of protein structures and complexes. Nucleic Acids Research, 46(W1), W296–W303. 10.1093/NAR/GKY427

