## Supplemental Information for "Decoding Sequence-Structure-Function-Evolution of basic Leucine Zippers of Aureochromes from Heterokont Algae"

### SI-1: Functions of different aureochrome homologs

| Name | Function | Reference |
|---|---|---|
| VfAureo1 | Promotes blue light induced branching | Takahashi et. al., 2007 PNAS |
| VfAureo2 | Suppresses the differentiation of branch primordium into sex organs | |
| SjAureo1 | Blue light dependent photo-morphogenesis during the growth and development of juvenile sporophyte | Deng et. al., 2014 Mar Biotechnol, Springer |
| NgAureo1 | Regulates lipid accumulation | Huang et. al., 2013 Biotechnol Lett, Springer |
| PtAureo1a | Controls G1 to S transition of cell cycle and regulates high light acclimation | Huysman et. al., 2013 Plant Cell |

### SI-2: List of all bZIPs used for Sequence comparison and molecular evolutionary analysis

**List of the selected bZIPs from the Opisthokont, Plant, Virus & Aureochrome for the construction of Phylogenetic Tree**

| UNIPROT ID | SEQUENCE OF THE bZIPs | NAME OF THE PROTEIN | Sub Group |
|---|---|---|---|
| | **Plant bZIPs** | | |
| Q9C5Q2 | VERRQKRMIKNRESAARSRARKQAYTHELEIKVSRLEEENEKLRRLKEVEKILPSEPPPDP | DPBF4_Arabidopsis thaliana | |
| Q9M7Q5 | VERRQKRMIKNRESAARSRARKQAYTLELEAEIESLKLVNQDLQKKQAEIMKTHNSELKE | ABF1_Arabidopsis thaliana | |
| Q9M7Q4 | VERRQRRMIKNRESAARSRARKQAYTVELEAEVAKLKEENDELQRKQARIMEMQKNQETE | ABF2 Arabidopsis thaliana | |
| Q9M7Q3 | IERRQKRMIKNRESAARSRARKQAYTMELEAEIAQLKELNEELQKKQVEIMEKQKNQLLEPL | ABF3 Arabidopsis thaliana | |
| Q9M7Q2 | IERRQRRMIKNRESAARSRARKQAYTLELEAEIEKLKKTNQELQKKQAEMVEMQKNELKE | ABF4_Arabidopsis thaliana | |
| Q9SJN0 | VERRQRRMIKNRESAARSRARKQAYTVELEAELNQLKEENAQLKHALAELERKRKQQYFESL | ABI5 Arabidopsis thaliana | |
| P42777 | QRQKRMIKNRESAARSRERKQAYQVELETLAAKLEEENEQLLKEIEESTKERYKKLMEVL | GBF4_Arabidopsis thaliana | |
| Q9LES3 | VERRQKRMIKNRESAARSRARKQAYTHELEIKVSRLEEENERLRKQKEVEKILPSVPPPDPK | DPBF3 Arabidopsis thaliana | |
| Q8RYD6 | MERRQRRMIKNRESAARSRARRQAYTVELELELNNLTEENTKLKEIVEENEKKRRQEIISRS | DPBF2 Arabidopsis thaliana | |
| Q84JK2 | GNRRHKRMIKNRESAARSRARKQAYTNELELEVAHLQAENARLKRQQDQLKMAAAIQQPKKNTL | FD Arabidopsis thaliana | A |
| Q7PCC6 | GDRRYKRMIKNRESAARSRARKQAYTNELELEIAHLQTENARLKIQQEQLKIAEATQNQVKK | FDP Arabidopsis thaliana | |
| Q5VQQ8 | MERRQKRMIKNRESAARSRARKQAYTNELENKVSRLEEENVRLKRQKESDYLHYTRSNLV | DPBF3_Oryza sativa | |
| Q6AVM5 | VERRQKRMIKNRESAARSRARKQAYTNELENKVLRLEEENERLKKQKELDEILNSAPPPE | AREB3like_Oryza sativa | |
| Q0JHF1 | AMQRQKRMIKNRESAARSRERKQAYIAELESLVTQLEEENAKMFKEQEEQHQKRLKELKEMV | BZIP12 Oryza sativa | |
| Q84P61 | AMQRQKRMIKNRESAARSRERKQAYIAELEAQVAELEEEHAQLLREQEEKNQKRLKEIKEQA | OSE2-like_Oryza sativa | |
| Q8RZ35 | VERRQRRMIKNRESAARSRARKQAYTVELEAELNYLKQENARLKEAEKTVLLTKKQMLVEKM | ABI5 Oryza sativa | |
| A0A3S5WM89 | AAQRQKRMIKNRESAARSRERKQAYQVELETLAAKLEEENEQLLKEIEESTKERYKKLMEVL | ABI5 Triticum aestivum | |
| Q67TQ5 | VERRQRRMIKNRESAARSRARKQAYTLELEAEVQKLKEMNKELERKQADIMEMQKNEVEEMI | Os09g0456200_Oryza sativa | |
| Q6ZDF3 | VERRQRRMIKNRESAARSRARKQAYTMELEAEVQKLKEQNMELQKKQEEIMEMQKNFFPEMQ | TRAB1_Oryza sativa | |

| | | | |
|---|---|---|---|
| Q8LIB3 | EAKRRARLVRNRESAHQSRQRKKQYVEELEGKVKVMQATIADLTARISCVTAENAALKQQL | BZIP60_Oryza sativa | B |
| Q6AU90 | DERRAARLMRNRESAQLSRQRKKRYVEELEEKVKSMHSVINDLNSRISFVVAENATLRQQL | BZIP39_Oryza sativa | |
| Q9SG86 | DKRKLIRQIRNRESAQLSRLRKKQQTEELERKVKSMNATIAELNGKIAYVMAENVALRQQM | BZIP28 Arabidopsis thaliana | |
| Q7X9A8 | DQRLQRRKQSNRESARRSRSRKAAHLNELEAQVSQLRVENSSLLRRLADVNQKYNDAAVDN | RISBZ2_Oryza sativa | C |
| Q6ZLB0 | EDKVKKRKESNRESARRSRSRKAARLKDLEEQVSLLRVENSSLLRRLADANQKYSAAAIDNRVL | RISBZ1_Oryza sativa | |
| B9DGI8 | NVKRVKRMLSNRESARRSRRRKQAHLSELETQVSQLRVENSKLMKGLTDVTQTFNDASVENRVL | BZIP63 Arabidopsis thaliana | |
| Q2QMJ8 | NAKRTRRMLSNRESARRSRKRKQAHLNDLESQVSQLRSENASLQKRLSDMTQKYKQSTTEYGNL | Os12g40920_Oryza sativa | |
| O22763 | DVKKSRRMLSNRESARRSRRRKQEQTSDLETQVNDLKGEHSSLLKQLSNMNHKYDEAAVGNRIL | BZIP10 Arabidopsis thaliana | |
| Q9M1G6 | DVKRARRMLSNRESARRSRRRKQEQMNEFDTQVGQLRAEHSTLINRLSDMNHKYDAAAVDNRIL | BZIP25 Arabidopsis thaliana | |
| Q6H500 | DVKRVRRMVSNRESARRSRKRKQAHLADLESQVDQLRGENASLFKQLTDANQQFTTSVTDNRIL | RISBZ4 Oryza sativa | |
| Q6ETX0 | DVKRMRRMVSNRESARRSRKRKQAHLADLETQVDQLRGENASLFKQLTDANQQFTTAVTDNR | RISBZ3 Oryza sativa | |
| Q654B3 | ETKRIRRMVSNRESARRSRRRKQAQLSELESQVEQLKGENSSLFKQLTESSQQFNTAVTDNRIL | RISBZ5 Oryza sativa | |
| Q9FUD3 | DLKRIRRMNSNRESAKRSRRRKQEYLVDLETQVDSLKGDNSTLYKQLIDATQQFRSAGTNNRVL | BZIP9 Arabidopsis thaliana | |
| P43273 | DQKTLRRLAQNREAARKSRLRKKAYVQQLENSRLKLTQLEQELQRARQQGVFISGTGDQAHST | TGA2 Arabidopsis thaliana | D |
| Q39234 | NDKMKRRLAQNREAARKSRLRKKAHVQQLEESRLKLSQLEQELVRARQQGLCVRNSSDTSYLG | TGA3 Arabidopsis thaliana | |
| Q39163 | DQKTLRRLAQNREAARKSRLRKKAYVQQLENSRLKLTQLEQELQRARQQGVFISSSGDQAHST | TGA5 Arabidopsis thaliana | |
| Q39140 | DQKTLRRLAQNREAARKSRLRKKAYVQQLENSRLKLTQLEQELQRARQQGVFISSSGDQAHST | TGA6 Arabidopsis thaliana | |
| Q9SX27 | DQRTLRRLAQNREAARKSRLRKKAYVQQLENSRIRLAQLEEELKRARQQGSLVERGVSADHTHL | PAN Arabidopsis thaliana | |
| Q39237 | PDKIQRRLAQNREAARKSRLRKKAYVQQLETSRLKLIQLEQELDRARQQGFYVGNGIDTNSL | TGA1 Arabidopsis thaliana | |
| Q39162 | PDKIQRRLAQNREAARKSRLRKKAYVQQLETSRLKLIHLEQELDRARQQGFYVGNGVDTNAL | TGA4 Arabidopsis thaliana | |
| Q93ZE2 | HDKMKRRLAQNREAARKSRLRKKAYVQQLEESRLKLSQLEQELEKVKQQGHLGPSGSINTG | TGA7 Arabidopsis thaliana | |
| Q5Z6N9 | DQKMQRRLAQNREAARKSRMRKKAYIQQLESSRSKLMHLEQELQRARQQGIFIATGGSGD | TGAL3 Oryza sativa | |
| Q2QXL0 | DPKIMRRLAQNREAARKSRLRKKAYIQQLESSKLRLAQMEQDLERARSQGLLLGGSPGGN | TGAL11_Oryza sativa | |
| Q6ZBS8 | RDKIQRRLAQNREAARKSRLRKKAYIQNLETSRMKLAHLEQEITRARQQSAYINRSSNPATL | TGAL10 _Oryza sativa | |
| Q6IVC3 | DQKVLRRLAQNREAARKSRLRKKAYVQQLESSKLKLASLEQEINKARQQGIYISSSGDQTH | TGA2.2 _Oryza sativa | |
| Q7X993 | DQKTIRRLAQNREAARKSRLRKKAYVQQLESSKLKLAQLEQELQKARQQGIFISSSGDQTH | TGA2.1 _Oryza sativa | |
| Q6IVC2 | DHKTLRRLAQNREAARKSRLRKKAYIQNLESSRLKLTQIEQELQRARQQGIFISTSSDQSH | TGAL1_Oryza sativa | |
| Q0JNL3 | DQKTLRRLAQNREAARKSRLRKKAYVQQLENSRLKLTQLEQELQRARQQGIFISSSVDQTH | TGA2.3_Oryza sativa | |

| | | | |
|---|---|---|---|
| F4IN23 | DPKRVKRILANRQSAQRSRVRKLQYISELERSVTSLQAEVSVLSPRVAFLDHQRLLLNVDN | BZIP34 Arabidopsis thaliana | E |
| Q9M2K4 | DPKRVKRILANRQSAQRSRVRKLQYISELERSVTSLQTEVSVLSPRVAFLDHQRLLLNVDN | BZIP61 Arabidopsis thaliana | |
| Q5JMK6 | DPKRVKRILANRQSAQRSRVRKLQYISELERSVTTLQNEVSVLSPRVAFLDQQRTILTVGNSHL | BZIP06_Oryza sativa | |
| Q5QNI5 | DPKRVKRILANRQSAQRSRVRKLQYISELERSVTSLQTEVSALSPRVAFLDHQRSLLNVDN | BZIP02 Oryza sativa | |
| Q8GTS1 | DSSNKKRLCGNREAVRKYREKKKARTAYLEDEVMRLQSLNEQFLRKLQSQEMVETELIRLRALL | BZIP24 _Arabidopsis thaliana | F |
| Q65X73 | DLTKTRRPLGNREAVRKYREKKKAHAAFLEEEVKKLRAANQQLLKRLQGHAALEAEVIRL | Os05g0495200 _Oryza sativa | |
| Q8VY76 | GKKGEKRPLGNREAVRKYREKKKAKAASLEDEVARLRAVNQQLVKRLQNQATLEAEVSRLKCLL | BZIP19 Arabidopsis thaliana | |
| Q5Z8P6 | NNASKKRPSGNRAAVRKYREKKKAHTASLEEEVVHLRALNQQLMKKLQNHATLEAEVSRLRCLL | Os06g0716800 protein Oryza sativa | |
| P42774 | RELKRQKRKQSNRESARRSRLRKQAECEQLQQRVESLSNENQSLRDELQRLSSECDKLKSENN | GBF1 Arabidopsis thaliana | G |
| P42775 | KEVKREKRKQSNRESARRSRLRKQAETEQLSVKVDALVAENMSLRSKLGQLNNESEKLRLENE | GBF2 Arabidopsis thaliana | |
| P42776 | RELKRERRKQSNRESARRSRLRKQAETEELARKVEALTAENMALRSELNQLNEKSDKLRGAN | GBF3 Arabidopsis thaliana | |
| Q40645 | ESKRERRKQSNRESARRSRLRKQAETEELARKVELLTAENTSLRREISRLTESSKKLRLENSAL | OSBZ8 Oryza sativa | |
| Q10PC3 | ELKRQKRKQSNRESARRSRLRKQAEWEEVANRADLLKQENSSLKEELKQLQEKCNSLTSENTTL | Os03g13614 Oryza sativa | |
| Q40625 | ELKRQRRKQSNRESARRSRLRKQAECEELAQRAEVLKQENTSLRDEVNRIRKEYDELLSKNSSL | osZIP-1a Oryza sativa | |
| O24646 | KENKRLKRLLRNRVSAQQARERKKAYLSELENRVKDLENKNSELEERLSTLQNENQMLRHILKN | HY5 Arabidopsis thaliana | H |
| Q8W191 | KEYRSLKRLLRNRVSAQQARERKKVYVSDLESRANELQNNNDQLEEKISTLTNENTMLRKMLIN | HYH Arabidopsis thaliana | |
| Q95M50 | KENKRLKRLLRNRVSAQQARERKKAYLIDLEARVKELETKNAELEERLSTLQNENQMLRHILKN | HY5 Solanum lycopersicum | |
| Q04088 | KRAKRIWANRQSAARSKERKTRYIFELERKVQTLQTEATTLSAQLTLLQRDTNGLTVENN | POSF21 Arabidopsis thaliana | I |
| Q2QXC3 | DPKRAKRILANRQSAARSKERKIKYTSELERKVQTLQTEATTLSAQLTLLQRDTSGLTAENREL | Os12g06520 Oryza sativa | |
| Q9MA75 | DPKRAKRILANRQSAARSKERKIRYTGELERKVQTLQNEATTLSAQVTMLQRGTSELNTENKHL | VIP1 Arabidopsis thaliana | |
| Q69IL4 | DPKRAKRIWANRQSAARSKERKMRYIAELERKVQTLQTEATTLSAQLALLQRDTSGLTTENSEL | RF2a Oryza sativa | |
| Q654P4 | RAKRILANRQSAARSKERKARYITELERKVQTLQTEATTLSAQLTLFQRDTTGLSAENA | RF2b Oryza sativa | |
| Q85A76 | DPKRVKRILANRQSAARSKERKMRYIQKLEHKVQVLQTEASTLSHSLKCLRRTPWTSPQNNEL | Vsf-1 Oryza sativa | |
| Q9SI15 | DERKRKRMLSNRESARRSRMRKQKHVDDLTAQINQLSNDNRQILNSLTVTSQLYMKIQAENSVL | BZIP2_Arabidopsis thaliana | S |
| O65683 | EQRKRKRMLSNRESARRSRMKKQKLLDDLTAQVNHLKKENTEIVTSVSITTQHYLTVEAENSVL | BZIP11 _Arabidopsis thaliana | |
| Q08418 | DERKRKRMLSNRESARRSRARKQQRLEELIAEAARLQAENARVEAQIGAYAGELSKVDGENAVL | lip19 _Oryza sativa | |
| Q4H4C4 | DHRREKRRLSNRESARRSRLRKQQHLDELVQEVARLQADNARVLARASEIAGQYARVEQENTVL | OsOBF1_Oryza sativa | |
| A9ZPJ3 | DNRREKRRLSNRESARRSRLRKQQHLDELVQEVARLKAENARVLARANDITGQFVRVDQENTVL | TaOBF1a _Triticum aestivum | |
| A9ZPJ4 | DNRREKRRLSNRESARRSRLRKQQHLDELVQEVARLKAENARVLARANDITGQFVRVDQENTVL | TaOBF1b _Triticum aestivum | |
| A9ZPJ5 | DNRREKRRLSNRESARRSRLRKQQHLDELVQEVARLKAENARVLARANDITGQFVRVDQENTVL | TaOBF1d_Triticum aestivum | |

| Opisthokont bZIPs | | | |
|---|---|---|---|
| Q01663 | EPSSKRKAQ**NRAAQRAFR**KRKEDHLKALETQVVTLKELHSSTTLENDQLRQKVRQLEEELRIL | PAP1_Schizosaccharomyces.pombe | |
| P19880 | ETKQKRTAQ**NRAAQRAFR**ERKERKMKELEKKVQSLESIQQQNEVEATFLRDQLITLVNELKK | YAP1_Saccharomyces.cerevisiae | |
| P38749 | DSKAKKKAQ**NRAAQKAFR**ERKEARMKELQDKLLESERNRQSLLKEIEELRKANTEINAENRLL | YAP3_ Saccharomyces.cerevisiae | PAP |
| Q03935 | GKTLRNTRRAAQ**NRTAQKAFR**QRKEKYIKNLEQKSKIFDDLLAENNNFKSLNDSLRNDNNILIAQ | YAP6_Saccharomyces.cerevisiae | |
| Q5AJU7 | KPIDTEPKSKRTAQ**NRAAQRAYR**ERKERKMKELEDKVRLLEDANVRALTETDFLRAQVDVLKNELAKYT | CAP1_Candida.albicans | |
| O54790 | KEEIIQLKQRRRTLK**NRGYAASCR**VKRVTQKEELEKQKAELQQEVEKLASENASMKLELDALRSKYEALQ | MafG_Mus.musculus | |
| P54841 | KDEVIRLKQKRRTLK**NRGYAQSCR**YKRVQQKHHLENEKTQLIQQVEQLKQEVSRLARERDAYKVKCEKLA | MafB_Mus.musculus | Maf |
| Q8NHW3 | KEEVIRLKQKRRTLK**NRGYAQSCR**FKRVQQRHILESEKCQLQSQVEQLKLEVGRLAKERDLYKEKYEKL | MafA_Homo.sapiens | |
| Q8N1L9 | EQQRQLKKQK**NRAAAQRSR**QKHTDKADALHQQHESLEKDNLALRKEIQSLQAELAWWSRTLHVH | BATF2_Homo.sapiens | |
| Q9NR55 | DDRKVRRREK**NRVAAQRSR**KKQTQKADKLHEEYESLEQENTMLRREIGKLTEELKHLTEALKEH | BATF3_Homo.sapiens | BATF |
| O35284 | DVRKVQRREK**NRIAAQKSR**QRQTQKADTLHLESEDLEKQNAALRKEIKQLTEELKYFTSVLSSH | BATF_Mus.musculus | |
| P17275 | QERIKVERKRLR**NRLAATKCR**KRKLERIARLEDKVKTLKAENAGLSSTAGLLREQVAQLKQKVM | JUN-B_Homo.sapiens | |
| P09450 | QERIKVERKRLR**NRLAATKCR**KRKLERIARLEDKVKTLKAENAGLSSAAGLLREQVAQLKQKVM | JUN-B_Mus.musculus | |
| K7PBE6 | RIKAERKRLR**NRLAATKCR**RRKLERIARLEEKVKVLKSDNAGLSNTASVLRDQVAQLKQEVLWH | jun-B_ Cyprinid herpesvirus1 | |
| P05412 | RIKAERKRMR**NRIAASKCR**KRKLERIARLEEKVKTLKAQNSELASTANMLREQVAQLKQKVMNH | JUN-C_Homo.sapiens | JUN |
| P52909 | QERIKAERKRLR**NRIAASKCR**KRKLERISRLEEKVKTLKSQNTELASTASLLREQVAQLKQKVL | JUN-D_ Rattus.norvegicus | |
| P15066 | QERIKAERKRLR**NRIAASKCR**KRKLERISRLEEKVKTLKSQNTELASTASLLREQVAQLKQKVL | JUN-D_Mus.musculus | |
| P17535 | QERIKAERKRLR**NRIAASKCR**KRKLERISRLEEKVKTLKSQNTELASTASLLREQVAQLKQKVL | JUN-D_Homo.sapiens | |
| P05411 | KAERKRMR**NRIAASKSR**KRKLERIARLEEKVKTLKAQNSELASTANMLREQVAQLKQKVMNH | JUN_Avian.sarcoma.virus | |
| Q5U0J5 | EAARKREVRLMK**NREAARECR**RKKKEYVKCLENRVAVLENQNKTLIEELKALKDLYCHKSD | CREB1_Homo.sapiens | |
| Q01147 | EAARKREVRLMK**NREAARECR**RKKKEYVKCLENRVAVLENQNKTLIEELKALKDLYCHKSD | CREB1_Mus.musculus | CREB |
| P18846 | DPQLKREIRLMK**NREAARECR**RKKKEYVKCLENRVAVLENQNKTLIEELKTLKDLYSNKSD | ATF1_Homo.sapiens | |
| P15336 | KRRKFLER**NRAAASRCR**QKRKVWVQSLEKKAEDLSSLNGQLQSEVTLLRNEVAQLKQLLLA | ATF2_Homo.sapiens | ATF2 |
| O93602 | DEKRRKFLER**NRAAASRCR**QKRKVWVQSLEKKAEDLSSLNGQLQNEVTLLRNEVAQLKQLLLAH | ATF2_ Gallus.gallus | |
| P18850 | VLRRQQRMIK**NRESACQSR**KKKKEYMLGLEARLKAALSENEQLKKENGTLKRQLDEVVSENQRL | ATF6A_Homo.sapiens | ATF6 |
| Q99941 | LLKRQQRMIK**NRESACQSR**RKKKEYLQGLEARLQAVLADNQQLRRENAALRRRLEALLAENSEL | ATF6B_ Homo.sapiens | |
| Q9XZS8 | EDRRRRRRER**NKIAATKCR**MKKRERTQNLIKESEVLDTQNVELKNQVRQLETERQKLVDMLKSH | ATF3_Drosophila.melanogaster | |
| P29596 | DERKRRRRER**NKIAAAKCR**NKKKEKTECLQKESEKLESVNAELKAQIEELKNEKQHLIYMLNL | ATF3_ Rattus norvegicus | ATF3 |
| P18847 | DERKKRRRER**NKIAAAKCR**NKKKEKTECLQKESEKLESVNAELKAQIEELKNEKQHLIYMLNLH | ATF3_Homo.sapiens | |
| O43889 | ILKRVRRKIR**NKRSAQESR**RKKKVYVGGLESRVLKYTAQNMELQNKVQLLEEQNLSLLDQLRKL | CREB3_Homo.sapiens | |
| F1P542 | ERLLKKVRRKIR**NKQSAQDSR**RRKKIYVDGLENRVAACTAQNHELQKKVQLLQKQNMSLLEQLR | CREB3L4_Gallus.gallus | |
| Q70SY1 | KALKKIRRKIK**NKISAQESR**RKKKEYMDSLEKKVESCSTENLELRKKVEVLENTNRTLLQQLQKLQTL | CREB3L2 or BBF2H7_Homo.sapiens | CREB3 |
| Q8BH52 | KALKKIRRKIKN**NKISAQESR**RKKKEYMDSLEKKVESCSTENLELRKKVEVLENTNRTLLQQLQKLQTL | CREB3L2 or BBF2H7_Mus.musculus | |
| Q9Z125 | ALKRVRRKIK**NKISAQESR**RKKKEYVECLEKKVETYTSENNELWKKVETLETANRTLLQQLQKL | CREB3L1/oasis_Mus.musculus | |
| Q96BA8 | ALKRVRRKIK**NKISAQESR**RKKKEYVECLEKKVETFTSENNELWKKVETLENANRTLLQQLQKL | CREB3L1/oasis_Homo.sapiens | |
| P18848 | EKLDKKLKKMEQ**NKTAATRYR**QKKRAEQEALTGECKELEKKNEALKERADSLAKEIQYLKDLIEEV | ATF4_Homo.sapiens | |
| Q06507 | EKLDKKLKKMEQ**NKTAATRYR**QKKRAEQEALTGECKELEKKNEALKEKADSLAKEIQYLKDLIEEV | ATF4_Mus.musculus | CREB2 |
| Q6NW59 | VEKKLKKMEQ**NKTAATRYR**QKKRVEQESLNSECSELEKKNRELSEKADSLSREIQYLRDLLEEM | ATF4_Danio.rerio | |
| Q16946 | HLDKKDRKKLQ**NKNAAIRYR**MKKKGEAQGIKGEEQELEELNTKLKTKVDDLQREIKYMKNLMEDV | ApCREB2_Aplysia californica | |
| P01100 | EKRRIRRER**NKMAAAKCR**NRRRELTDTLQAETDQLEDEKSALQTEIANLLKEKEKLEFILAAH | c-fos_Homo.sapiens | |
| P01101 | EKRRIRRER**NKMAAAKCR**NRRRELTDTLQAETDQLEDEKSALQTEIANLLKEKEKLEFILAAH | C-FOS_Mus.musculus | |
| P11939 | EEKRRIRRER**NKMAAAKCR**NRRRELTDTLQAETDQLEEEKSALQAEIANLLKEKEKLEFILAAH | c-Fos _Gallus.gallus | FOS |
| P53539 | EKRRVRRER**NKLAAAKCR**NRRRELTDRLQAETDQLEEEKAELESEIAELQKEKERLEFVLVAH | FOS-B_Homo.sapiens | |
| D3ZLB7 | EEKRRVRRER**NKLAAAKCR**NRRRELTDRLQAETDQLEEEKAELESEIAELQKEKERLEFVLVAH | FOSB_Rattus.norvegicus | |
| P15407 | EERRRVRRER**NKLAAAKCR**NRRKELTDFLQAETDKLEDEKSGLQREIEELQKQKERLELVLEAH | FRA1_Homo.sapiens | |
| P15408 | EEKRRIRRER**NKLAAAKCR**NRRRELTEKLQAETEELEEEKSGLQKEIAELQKEKEKLEFMLVAH | FRA2_Homo.sapiens | FRA |
| P10158 | ERRRVRRER**NKLAAAKCR**NRRKELTDFLQAETDKLEDEKSGLQREIEELQKQKERLELVLEAH | FRA1_Rattus.norvegicus | |
| P47930 | EEKRRIRRER**NKLAAAKCR**NRRRELTEKLQAETEELEEEKSGLQKEIAELQKEKEKLEFMLVAH | FRA2_Mus.musculus | |

| | | | |
|---|---|---|---|
| Q9Y4A8 | LIRDIRRRGK**NKVAAQNCR**KRKLDIILNLEDDVCNLQAKKETLKREQAQCNKAINIMKQKLHDL | NRF3_Homo.sapiens | CNC-Bzip |
| P20482 | LIRDIRRRGK**NKVAAQNCR**KRKLDQILTLEDEVNAVVKRKTQLNQDRDHLESERKRISNKFAML | CNC_Drosophila melanogaster | |
| Q00322 | DRGSPEYRQRRER**NNIAVRKSR**DKAKRRNQEMQQKLVELSAENEKLHQRVEQLTRDLAGLRQFFK | CRP3 or CEBPD_Mus.musculus | C/EBP |
| P49716 | DRGSPEYRQRRER**NNIAVRKSR**DKAKRRNQEMQQKLVELSAENEKLHQRVEQLTRDLAGLRQFFKQL | C/EBPD_Homo.sapiens | |
| P05554 | KNSNEYRVRRER**NNIAVRKSR**DKAKQRNVETQQKVLELTSDNDRLRKRVEQLSRELDTLRG | C/EBPA_Rattus.norvegicus | |
| P49715 | KNSNEYRVRRER**NNIAVRKSR**DKAKQRNVETQQKVLELTSDNDRLRKRVEQLSRELDTLRG | C/EBPA_Homo.sapiens | |
| P28033 | KKTVDKLSDEYKMRRER**NNIAVRKSR**DKAKMRNLETQHKVLELTAENERLQKKVEQLSRELSTLRNLFKQL | C/EBPB_Mus.musculus | |
| P17676 | KKTVDKHSDEYKIRRER**NNIAVRKSR**DKAKMRNLETQHKVLELTAENERLQKKVEQLSRELSTLRNLFKQL | C/EBPB_Homo.sapiens | |
| Q18909 | EPTYKLKRAR**NNDAVRKSR**KKAKELQDKKEAEHDKMKRRIAELEGLLQSERDARRRDQDTLEQLL | CEBP1_Caenorhabditis.elegans | |
| Q10586 | DEKYWSRRYK**NNEAAKRSR**DARRLKENQISVRAAFLEKENALLRQEVVAVRQELSHYRAVLSRY | DBP_Homo.sapiens | PAR |
| P16443 | DEKYWSRRYK**NNEAAKRSR**DARRLKENQISVRAAFLEKENALLRQEVVAVRQELSHYRAVLSRY | DBP_Rattus.norvegicus | |
| Q16534 | KDDKYWARRRK**NNMAAKRSR**DARRLKENQIAIRASFLEKENSALRQEVADLRKELGKCKNILAKY | HLF_Homo.sapiens | |
| Q64709 | KDDKYWARRRK**NNMAAKRSR**DARRLKENQIAIRASFLEKENSALRQEVADLRKELGKCKNILAKY | HLF_Rattus.norvegicus | |
| Q10587 | DEKYWTRRKK**NNVAAKRSR**DARRLKENQITIRAAFLEKENTALRTEVAELRKEVGKCKTIVSKY | TEF_Homo.sapiens | |
| O57673 | DDKYWQRRKK**NNVAAKRSR**DARRLKENQITVRAAFLERENSALRQEVAELRKDFGRCKNTVARY | TEF_Danio.rario | |
| P41224 | QKDEKYWTRRKK**NNVAAKRSR**DARRLKENQITIRAAFLEKENTALRTEVAELRKEVGKCKTIVSKY | TEF_Rattus.norvegicus | |
| Q16649 | KDAMYWEKRRK**NNEAAKRSR**EKRRLNDLVLENKLIALGEENATLKAELLSLKLKFGLISSTAYAQ | E4BP4_Homo.sapiens | E4BP4 |
| E2E3F5 | DNLYWERRRK**NNEAAKRSR**EKRRLNDMVLENKLMALGEENASLKAELLSLKLRFGLVSSAAYAQ | E4BP4_ Danio rerio | |
| Q90Z72 | DAMYWEKRRK**NNEAAKRSR**EKRRLNDLVLENKLIALGEENATLKAELLSLKLKFGLISSASYAQ | E4BP4_Gallus.gallus | |
| O08750 | KKDAMYWEKRRK**NNEAAKRSR**EKRRLNDLVLENKLIALGEENATLKAELLSLKLKFGLISSTAYAQ | E4BP4_Mus.musculus | |
| P03069 | SSDPAALKRAR**NTEAARRSR**ARKLQRMKQLEDKVEELLSKNYHLENEVARLKKLVSDPAALK | GCN4_Sachharomyces.cerevisiae | GCN4 |

| | | | |
|---|---|---|---|
| | **Aureochrome bZIPs** | | |
| A8QW55 | EAQKVERRER**NREHAKRSR**VRKKFLLESLQQSVNELNHENNCLKESIREHLGPRGDSL | VfAureo1 | Type-1 |
| V5RGX6 | EAQRVERRER**NREHAKRSR**MRKKFMLESLQAQMLALRKENLRLRQLVATKLPDKADTIL | NgAureo1 | |
| A0A1L4A1R0 | KIAEKIRKQR**NKEHAKRSR**VRKKFLVDSLQQSIDLLEKENQKLRSCLSANLGPEAQSII | HaAureo1 | |
| U3M7T2 | EQQKLERRER**NREHAKRSR**IRKKFMLECLQEQLLAMRKQNMALRQVVKEHMPNEAATVF | SjAureo1 | |
| C5NSW6 | EEQKIERRER**NREHAKRSR**VRKKFLLESLQHSVRALEEENEKLRNAIRENLQGEAEQLL | OdAureo1 | |
| D7FMH8 | EEQRIERRER**NREHAKRSR**VRKKFLLDSLQRSVDAIQAENESLKGSIVGSLGERGREL | EsAureo1 | |
| C5NSW4 | EEQRNERRER**NREHAKRSR**VRKKFLLDSLQRSVDALQAENDSLKGSIVGSLGERGREL | FeAureo1 | |
| C5NSW8 | STADQMRKQR**NKEHAKRSR**IRKKMLLDSLQKSIDLLEKENLKLRNTISNALGDEAKAL | CaAUREO1 | |
| K0SY93 | EKKSMDRRER**NREHAKRSR**IRKKFLLESLQQSVSLLKEENGKLKNAIRTHLGEKEAEAL | ToAureo1 | |
| B7G9J2 | QAQIDRRRER**NRILARRTR**LRKKFFFESLQKEIMDLQRENLVLKELVK | PtAureo1 | |
| F0YIK8 | EQQKDERRER**NREHAKRSR**VRKKFLLEPDYSLVKALQTAQQNFVITDPSL | AaAUREO1 | |
| W7U8A5 | HEQMERRRER**NRILARRTR**LRKKFIFESLQKQVMDLKRQNSRLKSIVKDKMADQASEVL | NgAureo2 | Type-2 |
| A0A1L4A1Q6 | EEQVERRRER**NRILARKTR**LRKKLFFEALQRRVTNLKTENELLRGVAQRRLGDADRRAAL | HaAureo2 | |
| D8LE91 | EEQAKKRRER**NRVLARRTR**LRKKFFFQSLQQQVARLQRENERLKGIVTTRCPDSVGEIL | EsAureo2 | |
| C5NSW5 | EEQAKKRRER**NRVLARRTR**LRKKFFFQSLQQQVNDLQYENERLKGIINTRCANNSAEI | FeAureo2 | |
| C5NSW7 | LTSKEKRKER**NKLLARKSR**MKLKADLENLKAKLMYLMKENESLRSQLYRVSTPPVSAEAL | OdAureo2 | |
| A8QW56 | EEQQQKRRER**NKVLARKTR**LRKKFLFQGLRNQVMSLYQENLRLKEIVKNHCGINSKKIL | VfAureo2 | |
| X2CR90 | EQQRLDRRER**NREHAKRSR**VRKKFLLESLQKSVTALQEENEKLRGAIRANLGADEAKEL | SjAureo2 | |
| B7FWL4 | EQQKLERRER**NREHAKRSR**LRKKFLLESLQEQIHGLEEQLDGLK | PtAureo2 | |
| A0A1L4A1R1 | EEQRLERRER**NREHAKRSR**IRKRFMLESLQEQFMGLQRENMALRQIIKDKIPARADSI | HaAureo3 | Type-3 |
| D8LT99 | EQQRLDRRER**NREHAKRSR**VRKKFLLESLQKSVTSLQEENEKLRGAIRSNLGPEEAKEL | EsAureo3 | |
| W7TTA1 | EEQKVERRER**NREHAKRSR**VRKKFLLESLQKSVNALQEENDKLRGAIRSHLKEGADDLL | NgAureo3 | |
| D7FRW8 | EQQKLERRER**NREHAKRSR**IRKKFMLECLQEQLLAMRKQNLALRQVVKEHMPDEASTV | EsAureo4 | Type-4 |
| W7U992 | EAQRVERRER**NREHAKRSR**MRKKFMLESLQAQMLALRKENLRLRQLVATKLPDKADTIL | NgAureo4 | |
| A0A1L4A1Q2 | MRKER**NKEHAKRSR**TRKKFLLDSFQYALDVLTKENEALRKNITASLGEEEAEREFAAFAEL | HaAureo4 | |
| A0A126X0V7 | GSELTDKRRR**NREHAKRSR**LRKRVRLGGLEEMVLGLRRENVRLRRIVKKGIPERADSILRL | SjAureo5 | Type-5 |

| | | | |
|---|---|---|---|
| | **Viral bZIPs** | | |
| A0A059ZXS9 | MLEIKRYK**NRVASRKCR**AKFKQLLQHYREVAAAKSSENDRLRLLLKQMCPSLDVDSIIPRTPD | ZEBRA (a variant of bZIP)_Epstein barr virus | Zta |
| A0A0X8Z9S8 | LEIKRYK**NRVASRKSR**AKFKQLLQHYREVAAAKSSENDRLRLLLKQMCPSLDVDSIIPRTPD | Epstein-Barr virus Zta DNA binding domain homodimer_Human gammaherpes virus 4 | |
| K7PBE6 | RIKAERKRLR**NRLAATKCR**RRKLERIARLEEKVKVLKSDNAGLSNTASVLRDQVAQLKQEVLWH | jun-B_ Cyprinid herpesvirus1 | Jun |
| P05411 | KAERKRMR**NRIAASKSR**KRKLERIARLEEKVKTLKAQNSELASTANMLREQVAQLKQKVMNH | JUN_Avian Sarcoma virus | |

### SI-3: Residue composition of the N-terminal basic region signature motif of all bZIPs

**ANIMAL bZIP subgroup**

```
PAP-   NRAAQKAFR
Maf-   NRGYAQSCR
BATF-  NRVAAQRSR
JUN-   NRIAASKCR
CREB-  NREAARECR
CREB3-NKISAQESR
CREB2-NKTAATRYR
FOS-   NKMAAAKCR
FRA-   NKLAAAKCR
C/EBP-NNIAVRKSR
PAR-   NNMAAKRSR
E4BP4-NNEAAKRSR
```

**AUREOCHROME bZIP**

Aureo1,3,4– NREHAKRSR
SjAureo2 -NREHAKRSR
HaAureo1 &
CaAureo1- NKEHAKRSR
VfAureo2- NKVLARKTR
EsAureo2 &
FeAureo2- NRVLARRTR
OdAureo2- NKLLARKSR
HaAu2 &
NgAu2- NRILARKTR

**PLANT bZIP subgroup**

```
A- NRESAARSR
B- NRESAQLSR
C- NRESARRSR
D- NREAARKSR
E- NRQSAQRSR
F- NREAVRKYR
G- NRESARRSR
H- NRVSAQQAR
I- NRQSAARSK
S- NRESARRSR
```

### SI-4: Graphs obtained from the Eigen-vector centrality of network constructed upon eight bZIP-DNA co-crystal structures of PDB

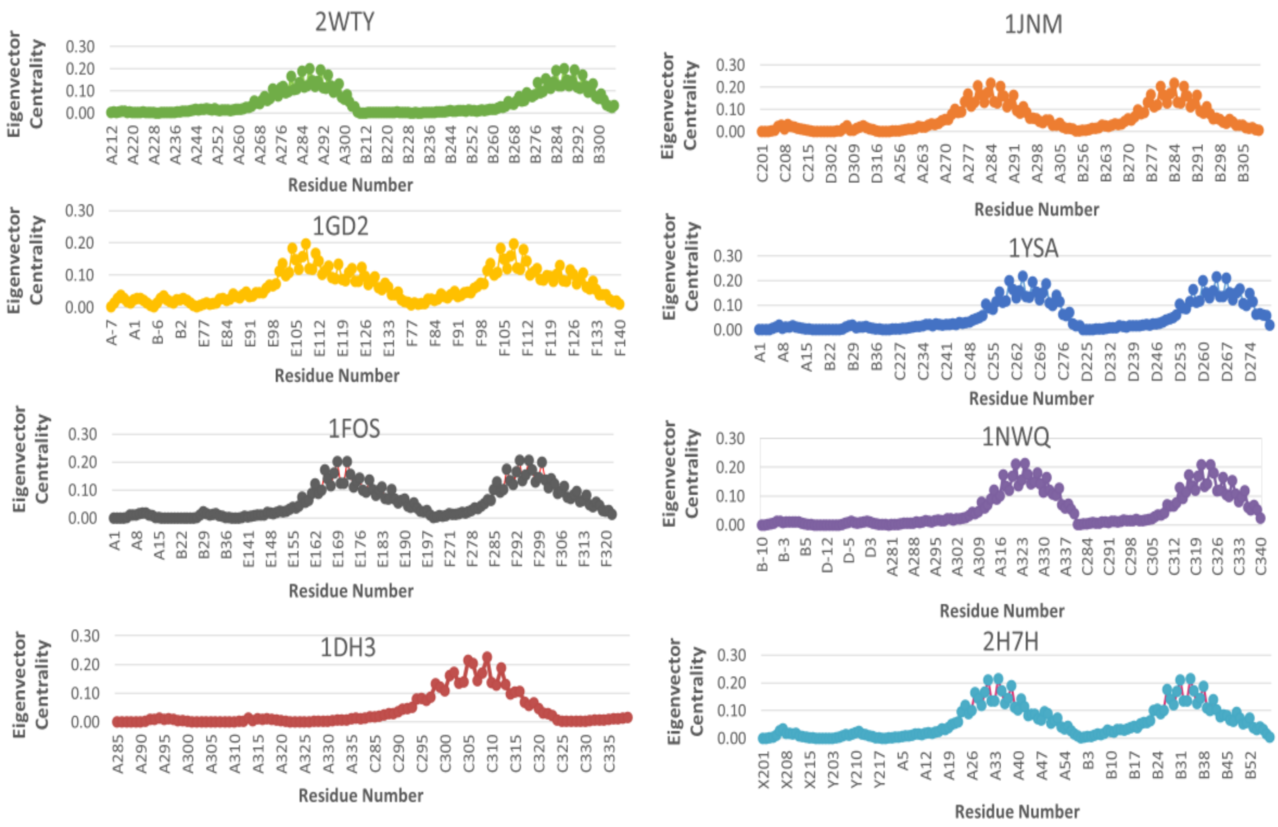

### SI-5: Hydrogen bonding details of the bZIP-DNA co-crystal structures

**Analysis of H-bonding interactions of Co-crystal structures of bZIP-DNA**

| PDB ID | Description | Chain No. | Basic region Residues | Hydrogen bonding details | Chain No. | Interacting DNA bases | Reference |
|---|---|---|---|---|---|---|---|
| **1GD2** | Pap with DNA | E | KRKAQNRAAQRAFRKRK | DA 4.A N6 ASN 86.E OD1<br>ARG 82.E NH2 DG -5.B O6<br>LYS 83.E NZ DA 3.A OP2<br>GLN 85.E NE2 DG -6.B OP2<br>ARG 87.E NH1 DT 2.A OP2<br>GLN 90.E NE2 DT -3.B O4<br>ARG 96.E NH2 DT -4.B OP2 | A<br>B | AGGTTACGTAACC<br>AGGTTACGTAACC | Fujii *et al*., 2000 |
| | | F | KRKAQNRAAQRAFRKRK | ARG`82.F NH2 DG`-6.A C8<br>DA 4.B N6 ASN 86.F OD1<br>LYS 81.F NZ DA 3.D N3<br>LYS 83.F NZ DA 3.B OP2<br>GLN 85.F NE2 DG -6.A OP2<br>ARG 87.F NH1 DT 2.B OP2<br>ARG 87.F NH2 DG 1.C OP1<br>GLN 90.F NE2 DT -3.A O4<br>ARG`96.F NH1 DT`-4.A OP2 | A<br>B<br>C<br>D | AGGTTACGTAACC<br>AGGTTACGTAACC<br>AGGTTACGTAACC<br>AGGTTACGTAACC | |
| | | G | KRKAQNRAAQRAFRKRK | ARG`94.G NH2 DG`1.C N7<br>DA 4.C N6 ASN 86.G OD1<br>LYS 81.G NZ DA 3.A N3<br>LYS 83.G NZ DA 3.C OP2<br>GLN 85.G NE2 DG -6.D OP2<br>ARG 87.G NH1 DT 2.C OP2<br>ARG 87.G NH2 DG 1.B OP1<br>GLN 90.G NE2 DT -3.D O4<br>ARG 96.G NH2 DT -4.D OP2<br>ARG`82.G NH2 DG`-6.D N7 | A<br>B<br>C<br>D | AGGTTACGTAACC<br>AGGTTACGTAACC<br>AGGTTACGTAACC<br>AGGTTACGTAACC | |
| | | H | KRKAQNRAAQRAFRKRK | DA 4.D N6 ASN 86.H OD1<br>ARG 82.H NH2 DG -5.C O6<br>LYS 83.H NZ DA 3.D OP2<br>GLN 85.H NE2 DG -6.C OP2<br>ARG 87.H NE DT 2.D OP2<br>GLN 90.H NE2 DT -3.C O4<br>ARG 94.H NH2 DG 1.D N7<br>ARG 96.H NH2 DT -4.C OP2 | C<br>D | AGGTTACGTAACC<br>AGGTTACGTAACC | |

| PDB ID | Description | Chain No. | Basic region Residues | Hydrogen bonding details | Chain No. | Interacting DNA bases | Reference |
|---|---|---|---|---|---|---|---|
| 1FOS | C-FOS:C-JUN hetero-dimer with DNA | F C-JUN | KAERKRMRNRIAASKSRKRK | ARG 268.F NE DC 34.B OP2<br>ASN 271.F ND2 DT 7.A O4<br>SER`278.F OG DT`7.A OP2<br>ARG`279.F NH2 DC`32.B N4 | A<br>B | AATGGATGAGTCATAGGAGA<br>TTCTCCTATGACTCATCCAT | Glover & Harrison, 1995 |
| | | H C-JUN | KAERKRMRNRIAASKSRKRK | ARG 268.H NE DT 11.C OP1<br>ASN 271.H ND2 DT 29.D O4<br>LYS 277.H NZ DA 28.D OP1<br>SER 278.H OG DT 29.D OP2<br>ARG 279.H NH1 DG 10.C N7<br>ARG 279.H NH2 DG 10.C N7<br>ARG 279.H NH2 DG 10.C O6<br>LYS 282.H NZ DG 30.D OP2 | C<br>D | AATGGATGAGTCATAGGAGA<br>TTCTCCTATGACTCATCCAT | |
| | | E C-FOS | KRRIRRERNKMAAAKSRNRRE | DC 12.A N4 ASN 147.E<br>ASN 147.E ND2 DT 29.B O4<br>LYS 153.E NZ DA 28.B OP1<br>SER 154.E OG DT 29.B OP2<br>ARG 155.E NH2 DG 10.A N7<br>ARG 158.E NH2 DG 30.B OP2 | A<br>B | AATGGATGAGTCATAGGAGA<br>TTCTCCTATGACTCATCCAT | |
| | | G C-FOS | KRRIRRERNKMAAAKSRNRRE | DC 34.D N4 ASN 147.G OD1<br>ASN 147.G ND2 DT 7.C O4<br>LYS 148.G NZ DT 33.D OP2<br>ARG`155.G NH2 DC`32.D C5<br>ARG`158.G NH2 DG`8.C OP2 | C<br>D | AATGGATGAGTCATAGGAGA<br>TTCTCCTATGACTCATCCAT | |

| | | | | | | | |
|---|---|---|---|---|---|---|---|
| **3A5T** | MafG with DNA | A | KQRRRTLKNRGYAASCRVKR | ARG 57.A NH1 DG 3.C N7<br>ARG 57.A NH2 DG 3.C O6<br>ARG 57.A NH2 DT 12.D O4<br>ARG 62.A NE DT 9.D OP2<br>ARG 62.A NH2 DT 9.D OP1<br>TYR 64.A OH DG 3.C OP2<br>CYS 68.A SG DT 5.C OP2<br>ARG 69.A NE DA 7.D OP2<br>LYS 71.A NZ DT 5.C OP1<br>DA 4.C N6 ASN 61.A OD1 | C<br>D | CTGATGAGTCAGCAC<br>GTGCTGACTCATCAG | Kurokawa *et al* ., 2009 |
| | | B | KQRRRTLKNRGYAASCRVKR | ARG 56.B NE DT 2.D OP2<br>ARG 56.B NH2 DT 2.D OP1<br>ARG 56.B NH2 DT 2.D OP2<br>ARG 57.B NH1 DG 3.D N7<br>ARG 57.B NH2 DG 3.D O6<br>LYS 60.B NZ DG 3.D OP2<br>ARG 62.B NE DT 9.C OP2<br>ARG 62.B NH2 DT 9.C OP1<br>TYR 64.B OH DG 3.D OP2<br>CYS 68.B SG DT 5.D OP2<br>ARG 69.B NE DA 7.C OP2<br>ARG 69.B NH2 DG 8.C OP2<br>LYS 71.B NZ DT 5.D OP1 | C<br>D | CTGATGAGTCAGCAC<br>GTGCTGACTCATCAG | |
| **2WTY** | MAFB with T-MARE | A | KRRTLKNRGYAQSCRYKRVQ | LYS 240.A NZ DT 3.D OP2<br>ARG 244.A NH1 DG 5.D N7<br>ARG 244.A NH2 DG 5.D O6<br>THR 245.A OG1 DC 12.C OP1<br>ARG 249.A NE DT 11.C OP1<br>TYR 251.A OH DG 5.D OP1<br>LYS 258.A NZ DT 7.D OP2<br>DC 6.D N4 ASN 248.A OD1 | C<br>D | AATTGCTGACTCAGCAAAT<br>ATTTGCTGAGTCAGCAATT | Pogenberg *et al* ., 2014 |
| | | B | KRRTLKNRGYAQSCRYKRVQ | ARG 244.B NH1 DG 5.C N7<br>ARG 244.B NH2 DG 5.C N7<br>ARG 244.B NH2 DG 5.C O6<br>ARG 244.B NH2 DG 14.D O6<br>THR 245.B OG1 DC 12.D OP1<br>ARG 249.B NE DT 11.D OP1<br>ARG 249.B NH2 DT 11.D OP1<br>TYR 251.B OH DG 5.C OP1<br>GLN 253.B NE2 DG 10.D OP2<br>CYS 255.B SG DT 7.C OP1<br>ARG 256.B NH1 DA 9.D OP1<br>DC 6.C N4 ASN 248.B OD1<br>LYS`240.B NZ DT`3.C OP2 | C<br>D | AATTGCTGACTCAGCAAAT<br>ATTTGCTGAGTCAGCAATT | |
| **4AUW** | Homo-dimeric MAFB with C-MARE | A | KRRTLKNRGYAQSCRYKRVQ | LYS 240.A NZ DA 4.D OP1<br>ARG 243.A NH2 DT 5.D OP1<br>ARG 244.A NH2 DG 16.C O6<br>ARG 244.A NH2 DG 6.D O6<br>LYS 247.A NZ DT 5.D OP1<br>ARG 249.A NE DG 12.C OP1<br>ARG 249.A NH2 DG 12.C OP1<br>TYR 251.A OH DG 6.D OP2<br>ARG 259.A NH2 DG 9.D OP2<br>DC 7.D N4 ASN 248.A OD1<br>GLN`253.A OE1 DC`11.C OP2 | C<br>D | TAATTGCTGACGTCAGCATTA<br>ATAATGCTGACGTCAGCAATT | Textor *et al* ., 2007 |
| | | B | KRRTLKNRGYAQSCRYKRVQ | LYS 240.B NZ DT 4.C OP1<br>ARG 244.B NH2 DG 6.C N7<br>ARG 244.B NH2 DG 16.D O6<br>ARG 249.B NE DT 13.D OP2<br>ARG 249.B NH2 DT 13.D OP1<br>TYR 251.B OH DG 6.C OP2<br>GLN 253.B NE2 DC 11.D OP2<br>CYS 255.B SG DT 8.C O5'<br>ARG 256.B NH2 DG 12.D N7<br>ARG 259.B NH2 DG 9.C OP2<br>DC 7.C N4 ASN 248.B OD1<br>ARG`243.B NH1 DT`5.C OP2 | C<br>D | TAATTGCTGACGTCAGCATTA<br>ATAATGCTGACGTCAGCAATT | |
| | | E | KRRTLKNRGYAQSCRYKRVQ | LYS 240.E NZ DA 4.G OP1<br>ARG 243.E NE DT 5.G OP2<br>ARG 243.E NH2 DT 5.G OP1<br>ARG 244.E NH2 DG 6.G O6<br>ARG 249.E NH1 DT 13.H OP2<br>TYR 251.E OH DG 6.G OP2<br>GLN 253.E NE2 DC 11.H OP2<br>CYS 255.E SG DT 8.G O5'<br>ARG 256.E NH1 DG 12.H N7<br>DC 7.G N4 ASN 248.E OD1<br>LYS`258.E NZ DT`8.G OP1 | G<br>H | ATAATGCTGACGTCAGCAATT<br>TAATTGCTGACGTCAGCATTA | |
| | | F | KRRTLKNRGYAQSCRYKRVQ | ARG 243.F NH2 DT 5.H OP1<br>ARG 244.F NH1 DG 6.H N7<br>ARG 244.F NH2 DG 6.H O6<br>TYR 251.F OH DG 6.H OP2<br>GLN 253.F NE2 DC 11.G OP2<br>CYS 255.F SG DT 8.H O5'<br>CYS 255.F SG DT 8.H OP2<br>ARG 256.F NH1 DC 11.G OP2<br>ARG 256.F NH2 DG 12.G N7<br>LYS 258.F NZ DT 8.H OP1<br>ARG`249.F NH1 DT`13.G OP2 | G<br>H | ATAATGCTGACGTCAGCAATT<br>TAATTGCTGACGTCAGCATTA | |
| **4EOT** | MafA homo-dimer bound to MARE | A | KRRTLKNRGYAQSCRFKRVQ | ARG 259.A NE DT 5.C OP2<br>ARG 259.A NH2 DT 5.C OP1<br>ARG 260.A NH1 DG 6.C N7<br>ARG 260.A NH2 DG 6.C N7<br>ARG 260.A NH2 DG 6.C O6<br>THR 261.A OG1 DC 12.D OP2<br>ARG 265.A NE DT 11.D OP2<br>ARG 265.A NH2 DT 11.D OP1<br>TYR 267.A OH DG 6.C OP2<br>GLN 269.A NE2 DC 10.D OP2<br>CYS 271.A SG DT 8.C OP2<br>DC 7.C N4 ASN 264.A OD1 | C<br>D | CCGGTGCTGAGTCAGCAGG<br>CCCTGCTGACTCAGCACCG | Lu *et al* ., 2012 |
| | | B | KRRTLKNRGYAQSCRFKRVQ | ARG 260.B NH1 DG 5.D N7<br>ARG 260.B NH2 DG 15.C O6<br>ARG 260.B NH2 DG 5.D N7<br>ARG 260.B NH2 DG 5.D O6<br>THR 261.B OG1 DC 13.C OP2<br>ARG 265.B NE DT 12.C OP2<br>ARG 265.B NH2 DT 12.C OP2<br>TYR 267.B OH DG 5.D OP2<br>CYS 271.B SG DT 7.D OP2<br>ARG 272.B NH2.A DG 11.C O6<br>LYS 274.B NZ DT 7.D OP1<br>DC 6.D N4 ASN 264.B OD1<br>ARG`259.B NH2 DT`4.D OP1 | C<br>D | CCGGTGCTGAGTCAGCAGG<br>CCCTGCTGACTCAGCACCG | |

| | | | | | | | |
|---|---|---|---|---|---|---|---|
| **1JNM** | JUN with CRE DNA | A | KAERKRMRNRIAASKSRKRK | ARG 261.A NE DG 305.D OP2<br>ASN 262.A ND2 DT 307.D O4<br>ARG 263.A NE DG 211.C O5<br>ARG 263.A NE DG 211.C OP1<br>ARG 263.A NH2 DG 211.C OP1<br>SER 269.A OG DT 307.D OP2<br>DC 213.C N4 ASN 262.A OD1<br>ARG`270.A NH1 DC`210.C OP2 | C | CGTCGATGACGTCATCGACG | Kim *et al* . |
| | | | | | D | CGTCGATGACGTCATCGACG | |
| | | B | KAERKRMRNRIAASKSRKRK | ARG 261.B NE DG 205.C OP2<br>ASN 262.B ND2 DT 207.C O4<br>SER 269.B OG DT 207.C OP2<br>ARG 270.B NH2 DG 311.D O6<br>DC 313.D N4 ASN 262.B OD1<br>ARG`263.B CD DG`311.D OP2 | C | CGTCGATGACGTCATCGACG | |
| | | | | | D | CGTCGATGACGTCATCGACG | |
| **5T01** | C-Jun homo-dimer with methylated DNA | A | KAERKRMRNRIAASKSRKRK | ARG 259.A NH1 DT 11.C OP1<br>ARG 261.A NH2 DG 27.D OP2<br>ARG 263.A NE DT 11.C OP2<br>ARG 263.A NH2 DT 11.C OP1<br>ARG 270.A NE DA 9.C OP2<br>ARG 270.A NH2 DC 10.C OP2 | C | CTCCTATGACTCGTCCAT | Hong *et al* ., 2017 |
| | | | | | D | AATGGAXGAGTCATAGGAG | |
| | | B | KAERKRMRNRIAASKSRKRK | ARG 259.B NH1 DC 34.D OP2<br>ARG 261.B NE DT 5.C OP1<br>ARG 261.B NH2 DT 5.C OP1<br>ASN 262.B ND2 DT 7.C O4<br>SER 269.B OG DT 7.C OP2<br>ARG 270.B NH1 DG 32.D N7<br>ARG 270.B NH2 DG 32.D O6<br>ARG 272.B NH2 DT 7.C OP2<br>LYS 273.B NZ DG 8.C OP2<br>DC 34.D N4 ASN 262.B OD1<br>ARG 263.B CG DG 32.D OP2 | C | CTCCTATGACTCGTCCAT | |
| | | | | | D | AATGGAXGAGTCATAGGAG | |
| **2H7H** | JUN (Homo) with AP1 | A | KAERKRMRNRIAASKSRKRK | ARG 6.A NE.B DC 212.X OP2<br>ARG 6.A NH1.B DC 212.X OP2<br>ARG 8.A NE DG 205.Y OP2<br>ARG 8.A NH1 DC 204.Y OP2<br>ARG 8.A NH1 DG 205.Y OP2<br>ARG 10.A NH1 DT 211.X OP2<br>LYS 15.A NZ DA 206.Y OP1<br>ARG 17.A NE.B DA 209.X OP2<br>LYS 20.A NZ DG 208.Y OP2 | X | CGTCGATGACTCATCGACG | Kim *et al.* |
| | | | | | Y | CGTCGATGAGTCATCGACG | |
| | | B | KAERKRMRNRIAASKSRKRK | ARG 6.B NH1.A DT 211.Y OP1<br>ARG 8.B NE DG 205.X OP2<br>ARG 8.B NH2 DG 205.X OP2<br>ARG 10.B NE DT 211.Y OP2<br>LYS 15.B NZ DA 206.X OP1<br>SER 16.B OG DT 207.X OP2<br>ARG 17.B NE.B DA 209.Y OP2<br>ARG 17.B NH2.A DG 210.Y N7<br>LYS 20.B NZ DG 208.Y OP2 | X | CGTCGATGACTCATCGACG | |
| | | | | | Y | CGTCGATGAGTCATCGACG | |
| **1DH3** | CREB BZIP-CRE | A | VRLMKNREAARESRRKKKEY | ARG 286.A NH2 DC 3.B OP1<br>ARG 294.A NE DT 2.B OP2<br>ARG 298.A NH2 DG 1.B OP1<br>ARG 301.A NH1 DC -1.B OP2<br>ARG 301.A NH2 DG 1.B N7<br>DC 3.B N4 ASN 293.A OD1 | B | CCTTGGCTGACGTCAGCCAAG | Schumacher *et al* ., 2000 |
| | | C | VRLMKNREAARESRRKKKEY | ASN 293.C ND2 DT -4.B O4<br>LYS 305.C NZ DA -2.D OP1<br>DC 3.D N4 ASN 293.C OD1<br>ARG 294.C NH1 DG 1.D O3<br>ARG 298.C NH2 DG 1.D OP2<br>ARG 286.C NH2 DC 3.D OP2 | D | CCTTGGCTGACGTCAGCCAAG | |
| | | | | | B | CCTTGGCTGACGTCAGCCAAG | |

| | | | | | | | |
|---|---|---|---|---|---|---|---|
| **1NWQ** | C/EBP Alpha with DNA | A | RVRRERNNIAVRKSRDKAKQ | TYR 285.A OH DA 3.D OP2<br>ARG 286.A NH2 DC 2.D OP1<br>ARG 289.A NE DC 2.D OP2<br>ARG 289.A NH2 DC 2.D O5'<br>ARG 289.A NH2 DC 2.D OP2<br>ASN 293.A ND2 DC 2.D OP2<br>LYS 298.A NZ DA -5.B OP1<br>SER 299.A OG DT -4.B OP2<br>ARG 300.A NH1 DG 1.D O6<br>ARG 300.A NH2 DG 1.D O6<br>LYS 302.A NZ DT -4.B OP2<br>DA 3.D N6 ASN 292.A OD1 | B | TTCCTATTGCGCAATCCAGTT | Miller *et al*., 2003 |
| | | | | | D | AAACTGGATTGCGCAATAGGA | |
| | | C | RVRRERNNIAVRKSRDKAKQ | DA 3.B N6 ASN 292.C OD1<br>TYR 285.C OH DA 3.B OP2<br>ARG 289.C NE DC 2.B OP2<br>ARG 289.C NH1 DA 3.B N7<br>ASN 292.C ND2 DT -4.D O4<br>ASN 293.C ND2 DC 2.B OP2<br>SER 299.C OG DT -4.D OP2<br>ARG 300.C NH1 DG 1.B N7<br>ARG 300.C NH1 DG 1.B O6<br>ARG 300.C NH2 DG 1.B O6<br>LYS 304.C NZ DG -2.B OP1<br>ARG 286.A NH2 DC 2.B OP1<br>LYS 298.A NZ DA -5.D OP1 | B | TTCCTATTGCGCAATCCAGTT | |
| | | | | | D | AAACTGGATTGCGCAATAGGA | |
| **2 E43** | C/EBP-beta mutant bZIP homo-dimer bound to A-DNA | A | KIRRERNNIAVRKSRDKAK | TYR 274.A OH DA 12.C OP2<br>ARG 278.A NE DC 11.C OP2<br>ARG 278.A NH1 DA 12.C N7<br>ASN 281.A ND2 DT 105.D O4<br>ASN 282.A ND2 DC 11.C OP2<br>LYS 287.A NZ DA 104.D OP1<br>SER 288.A OG DT 105.D OP2<br>ARG 289.A NH1 DG 10.C O6<br>ARG 289.A NH2 DG 10.C N7<br>LYS 293.A NZ DG 8.C OP2<br>DA 12.C N6 ASN 281.A OD1<br>ARG 280.B NH2 DT 103.D OP1 | C | TAGGATTGCGCAATAT | Tahirov *et al.* |
| | | | | | D | AATATTGCGCAATCCT | |
| | | B | KIRRERNNIAVRKSRDKAK | TYR 274.B OH DA 111.D OP2<br>ARG 278.B NE DC 110.D OP2<br>ARG 278.B NH1 DA 111.D N7<br>ARG 280.B NE DG 4.C OP2<br>ASN 281.B ND2 DT 6.C O4<br>ASN 282.B ND2 DC 110.D OP2<br>LYS 287.B NZ DA 5.C OP1<br>SER 288.B OG DT 6.C O5'<br>SER 288.B OG DT 6.C OP2<br>LYS 291.B NZ DT 6.C OP1<br>DA 111.D N6 ASN 281.B OD1 | C | TAGGATTGCGCAATAT | |
| | | | | | D | AATATTGCGCAATCCT | |
| **1GTW** | C/EBP-beta bZip homodimer bound to a DNA | A | EYKIRRERNNIAVRKSRDKA | LYS 269.A NZ DA 11.C OP1<br>TYR 274.A OH DA 11.C OP2<br>ARG 278.A NE DC 10.C OP2<br>ARG 280.A NH1 DG 103.D OP2<br>ASN 281.A ND2 DT 106.D O4<br>ASN 282.A ND2 DC 10.C OP2<br>LYS 287.A NZ DA 105.D OP1<br>SER 288.A OG DT 106.D OP2<br>ARG 289.A NH1 DG 9.C O6<br>ARG 289.A NH2 DG 9.C N7<br>LYS 293.A NZ DG 7.C OP2<br>DA 11.C N6 ASN 281.A OD1 | C | AATGTGGCGCAATCCT | Tahirov *et al.* |
| | | | | | D | TAGGATTGCGCCACAT | |
| | | B | EYKIRRERNNIAVRKSRDKA | LYS 269.B NZ DC 112.D OP1<br>TYR 274.B OH DC 112.D OP2<br>ARG 278.B NE DC 111.D OP2<br>ARG 278.B NH2 DC 111.D OP2<br>ASN 281.B ND2 DT 5.C O4<br>ASN 282.B ND2 DC 111.D OP2<br>LYS 287.B NZ DG 4.C OP1<br>SER 288.B OG DT 5.C OP2<br>ARG 289.B NH1 DG 110.D O6<br>ARG 289.B NH2 DG 110.D O6<br>LYS 291.B NZ DT 5.C OP1<br>DC 112.D N4 ASN 281.B OD1 | C | AATGTGGCGCAATCCT | |
| | | | | | D | TAGGATTGCGCCACAT | |
| **1YSA** | GCN4 with DNA | C | KRARNTEAARRSRARK | DC 12.A N4 ASN 235.C OD1<br>ARG 234.C NE DG 27.B OP2<br>ARG 234.C NH2 DG 27.B OP2<br>ASN 235.C ND2 DT 29.B O4<br>THR 236.C OG1 DT 11.A OP2<br>ARG 241.C NH1 DA 28.B OP2<br>ARG 243.C NH1 DA 9.A OP2<br>ARG 245.C NH2 DT 29.B OP2 | A | TTCCTATGACTCATCCAGTT | Ellenberger *et al*., 1992 |
| | | | | | B | AAACTGGATGAGTCATAGGA | |
| | | D | AALKRARNTEAARRSRARK | DC 34.B N4 ASN 235.D OD1<br>ASN 235.D ND2 DT 7.A O4<br>THR 236.D OG1 DT 33.B OP2<br>ARG 241.D NH2 DA 6.A OP2<br>ARG 243.D NH1 DG 32.B O6<br>ARG 243.D NH2 DG 32.B N7<br>ARG 245.D NH2 DT 7.A OP2 | A | TTCCTATGACTCATCCAGTT | |
| | | | | | B | AAACTGGATGAGTCATAGGA | |

| | | | | | | | |
|---|---|---|---|---|---|---|---|
| 1GU4 | C/EBP BETA bound to DNA | A | KHSDEYKIRRERNNIAVRK SRDKAK | LYS 269.A NZ DA 12.C OP1<br>TYR 274.A OH DA 12.C OP2<br>ARG 278.A NE DC 11.C OP2<br>ARG 278.A NH1 DA 12.C N7<br>ASN 281.A ND2 DT 105.D O4<br>ASN 282.A ND2 DC 11.C OP2<br>LYS 287.A NZ DA 104.D OP1<br>SER 288.A OG DT 105.D OP2<br>ARG 289.A NH1 DG 10.C O6<br>ARG 289.A NH2 DG 10.C N7<br>LYS 293.A NZ DG 8.C OP2<br>DA 12.C N6 ASN 281.A OD1 | C<br><br>D | TAGGATTGCGCAATAT<br><br>AATATTGCGCAATCCT | Tahirov *et al.* |
| | | B | KHSDEYKIRRERNNIAVRK SRDKAK | LYS 269.B NZ DA 111.D OP1<br>TYR 274.B OH DA 111.D OP2<br>ARG 278.B NE DC 110.D OP2<br>ARG 278.B NH2 DC 110.D OP2<br>ASN 281.B ND2 DT 6.C O4<br>ASN 282.B ND2 DC 110.D OP2<br>LYS 287.B NZ DA 5.C OP1<br>SER 288.B OG DT 6.C OP2<br>ARG 289.B NH1 DG 8.C O6<br>ARG 289.B NH2 DG 8.C N7<br>LYS 291.B NZ DT 6.C OP1<br>DA 111.D N6 ASN 281.B OD1 | C<br><br>D | TAGGATTGCGCAATAT<br><br>AATATTGCGCAATCCT | |
| 1DGC | GCN4 with ATF/ CREB | A | KRARNTEAARRSRARK | ARG 240.A NH1 DC -1.B O5'<br>ARG 240.A NH2 DG 1.B OP2<br>ARG 243.A NH1 DC -1.B OP2<br>DC 3.B N4 ASN 235.A OD1<br>THR 236.A OG1 DG 1.B OP2 | B | TGGAGATGACGTCATCTCC | König & Richmond, 1993 |
| 2DGC | | A | KRARNTEAARRSRARK | THR 236.A OG1 DT 2.B OP2<br>ARG 240.A NH2 DG 1.B OP2<br>ARG 243.A NH2.A DG 1.B N7<br>ARG 243.A NH2.B DC -1.B OP2<br>DC 3.B N4 ASN 235.A OD1 | B | TGGAGATGACGTCATCTCC | |

| | | | | | | | |
|---|---|---|---|---|---|---|---|
| **Crystal structures of the bZIP Heterodimers** | | | | | | | |
| 2WT7 | MafB-cFos hetero-dimer with DNA | A | EKRRIRRERNKMAAAKCRNRRRE | ASN 147.A ND2 DT 4.D O4<br>LYS 153.A NZ DA 3.D OP1<br>CYS 154.A SG DT 4.D OP2<br>ARG 155.A NE DA 9.C OP2<br>ARG 159.A NH2.A DA 9.C OP2<br>DC 12.C N4 ASN 147.A OD1 | C<br><br>D | AATTGCTGACTCATAG<br><br>CTATGAGTCAGCAATT | Pogenberg *et al*., 2014 |
| | | B | )KRRTLKNRGYAQSCRYKRVQQKI | LYS 240.B NZ DT 3.C OP1<br>ARG 243.B NE DT 4.C OP2<br>ARG 243.B NH2 DT 4.C OP1<br>ARG 244.B NH1 DG 5.C N7<br>ARG 244.B NH2 DG 5.C O6<br>THR 245.B OG1 DC 9.D OP2<br>ARG 249.B NE DT 8.D OP2<br>ARG 249.B NH2 DT 8.D OP1<br>TYR 251.B OH DG 5.C OP2<br>GLN 253.B NE2 DG 7.D OP2<br>CYS 255.B SG DT 7.C OP2<br>ARG 256.B NH1 DG 7.D N7<br>ARG 256.B NH2 DG 7.D O6<br>LYS 258.B NZ DC 6.C OP1 | C<br><br>D | AATTGCTGACTCATAG<br><br>CTATGAGTCAGCAATT | |
| 5ZK1 | CRTC2-CREB-CRE | A | (REVRLMKNREAARESRRKKKEY\ | ARG 301.A NH1 DC -1.B OP2<br>LYS 305.A NZ DA -2.B OP1<br>DC 3.B N4 ASN 293.A OD1 | B | CTTGGCTGACGTCAGCCAAG | Song *et al*., 2018 |
| 1T2K | ATF-2 and Jun Bound To DNA | D (ATF 2) | RKFLERNRAAASRSRQKRKVWV( | ARG 343.D NE DA 3.E OP1<br>ARG 343.D NH2 DA 3.E OP1<br>ASN 344.D ND2 DT 5.E O4<br>ARG 345.D NE DT 26.F OP2<br>ARG 345.D NH2 DT 26.F OP2<br>ARG 352.D NH2 DG 25.F N7<br>LYS 354.D NZ DT 5.E OP1<br>DC 27.F N4 ASN 344.D OD1 | E<br><br>F | TAAATGACATAGGAAAACTGA<br><br>TCCTATGTCATTT | Panne *et al*., 2004 |

| | | | | | | | |
|---|---|---|---|---|---|---|---|
| 5VPE | FosB/JunD with cognate DNA | B<br>D-JUN | QERIKAERKRLRNRIAASKCRKR | ARG 275.B NH1 DC 12.E OP2<br>ARG 277.B NH1.B DG 5.F OP2<br>ASN 278.B ND2 DT 7.F O4<br>ARG 279.B NH1.B DT 11.E OP2<br>LYS 284.B NZ DG 6.F OP1<br>CYS 285.B SG DT 7.F OP2<br>ARG 286.B NH2.B DA 9.F N6<br>LYS 289.B NZ DG 8.F OP2<br>DC 12.E N4 ASN 278.B OD1<br>ARG`275.B HD3 DT`11.E O3'<br>ARG`288.B HH22 DT`7.F OP2 | E<br>F | CGTCGGTGACTCACCGACG<br>CGTCGGTGAGTCACCGACG | Yin *et al*., 2017 |
| | | D<br>D-JUN | QERIKAERKRLRNRIAASKCRKR | ARG 275.D NH1 DC 12.G OP2<br>ASN 278.D ND2 DT 7.H O4<br>ARG 279.D NE.B DT 11.G OP2<br>ARG 279.D NH2.B DT 11.G OP2<br>LYS 284.D NZ DG 6.H OP1<br>CYS 285.D SG DT 7.H OP2<br>ARG 286.D NH2.A DA 9.G OP2<br>ARG 286.D NH2.B DT 11.G O4<br>ARG 288.D NH2 DT 7.H OP2<br>LYS 289.D NZ DG 8.H OP2<br>DC 12.G N4 ASN 278.D OD1<br>ARG`277.D HG2 DG`19.E O3' | G<br>H | CGTCGGTGACTCACCGACG<br>CGTCGGTGAGTCACCGACG | |
| | | A | EEEKRRVRRERNKLAAAKCRNRR | ARG 161.A NH2 DG 6.E O6<br>ARG 164.A NH1.A DC 4.E OP1<br>ASN 165.A ND2 DT 7.E O4<br>LYS 171.A NZ DG 6.E OP1<br>CYS 172.A SG.A DT 7.E OP2<br>ARG 173.A NH1 DG 10.F N7<br>ARG 173.A NH2 DG 10.F O6<br>ARG 176.A NH2 DG 8.E OP2<br>DC 12.F N4 ASN 165.A OD1 | E<br>F | CGTCGGTGACTCACCGACG<br>CGTCGGTGAGTCACCGACG | |
| | | C | EEEKRRVRRERNKLAAAKCRNRR | ARG 161.C NH2 DG 6.G O6<br>ASN 165.C ND2 DT 7.G O4<br>LYS 171.C NZ DG 6.G OP1<br>CYS 172.C SG.A DT 7.G OP2<br>CYS 172.C SG.B DT 7.G OP2<br>ARG 173.C NH1 DG 10.H N7<br>ARG 173.C NH2 DG 10.H O6<br>ARG 176.C NH1 DG 8.G OP2<br>ARG 176.C NH2 DG 8.G OP1<br>ARG 177.C NH1 DA 9.H OP2<br>ARG 177.C NH2 DA 9.H OP2<br>DC 12.H N4 ASN 165.C OD1 | G<br>H | CGTCGGTGACTCACCGACG<br>CGTCGGTGAGTCACCGACG | |

| | | | | | | | |
|---|---|---|---|---|---|---|---|
| 5VPF | FosB/JunD with cognate DNA | B<br>(D-JUN) | QERIKAERKRLRNRIAASKCRKR | ASN 278.B ND2 DT 7.F O4<br>ARG 279.B NH1.A DT 11.E OP2<br>LYS 284.B NZ DG 6.F OP1<br>CYS 285.B SG DT 7.F OP2<br>ARG 288.B NH1.A DT 7.F OP2<br>ARG 288.B NH2.A DT 7.F OP2<br>LYS 289.B NZ DG 8.F OP2<br>DC 12.E N4 ASN 278.B OD1 | E<br>F | CGTCGGTGACTCACCGACG<br>CGTCGGTGAGTCACCGACG | Yin *et al*., 2017 |
| | | D<br>(D-JUN) | QERIKAERKRLRNRIAASKCRKR | ARG 277.D NE DG 5.H OP2<br>ASN 278.D ND2 DT 7.H O4<br>CYS 285.D SG DT 7.H OP2<br>ARG 286.D NH2.B DA 9.H N6<br>LYS 289.D NZ DG 8.H OP2<br>DC 12.G N4 ASN 278.D OD1 | G<br>H | CGTCGGTGACTCACCGACG<br>CGTCGGTGAGTCACCGACG | |
| | | A | EEEKRRVRRERNKLAAAKCRNRR | ARG 161.A NH1 DG 5.E N7<br>ARG 161.A NH2 DG 6.E O6<br>ARG 164.A NE DG 5.E OP2<br>ASN 165.A ND2 DT 7.E O4<br>CYS 172.A SG.A DT 7.E OP2<br>ARG 173.A NH1.A DG 10.F N7<br>ARG 173.A NH2.A DG 10.F O6<br>DC 12.F N4 ASN 165.A OD1 | E<br>F | CGTCGGTGACTCACCGACG<br>CGTCGGTGAGTCACCGACG | |
| | | C | EEEKRRVRRERNKLAAAKCRNRR | ARG 161.C NH2 DG 6.G O6<br>ASN 165.C ND2 DT 7.G O4<br>CYS 172.C SG DT 7.G OP2<br>ARG 173.C NH1.A DG 10.H N7<br>ARG 173.C NH2.A DG 10.H N7<br>ARG 173.C NH2.A DG 10.H O6<br>ARG 173.C NH2.B DA 9.H OP2<br>DC 12.H N4 ASN 165.C OD1 | G<br>H | CGTCGGTGACTCACCGACG<br>CGTCGGTGAGTCACCGACG | |

**** Residues that are involved in more than one interactions are underligned. If it is taking part in 3 interactions, then it is double underlined.**

### SI-6: Detailed polar interactions of aureochrome-bZIP with DNA

The N-terminal basic regions being the key determinants behind DNA binding specificity of bZIPs, they were first screened for polar including hydrogen bonding interactions with the DNA substrate. All interactions are color coded and presented in tabular format [SI-3]. In EsAureo1-DNA complex, Arg355 (NH2) forms hydrogen bond with dC’14 (OP2), while Arg360 (NH1) interacts with dG’12 (OP1). Asn359 interacts with N4 of dC’14 via hydrogen bonding, whereas Lys371 (NZ) engages with dA’10 (O5). His362 (NE2) interacts with dT’8 (O4). In EsAureo2, Arg190 (NH1), Arg193 (NH2), Asn194 (ND), and Arg199 (NH1) form polar contacts with dC’14 (OP2), dG’5 (OP2), dT’8 (O4) and dG’12 (OP1) respectively. In EsAureo3, His51 and Asn48 are in close proximity to multiple deoxy-ribonucleotides liked dG’6, dA’15 and dT’13. Additionally, Arg47 (NH1), Ser55 (OG), Arg54 (NE) and Arg56 (NH2) make polar contacts with dC’7 (OP2), dA’10 (N7), dG’9 (OP2) and dG’9 (N7) respectively. The residues from the basic region of EsAureo4 like Arg55 (NH2), Asn47 (ND2), Arg53 (NH1), Arg46 (NH1) and His50 (NE2) also show similar interaction pattern like in other Aureos. The respective interacting partner residues from the substrate are dG’9 (OP2), dG’16 (O6), dG’6 (OP2), dC’7 (OP2) and dT’8 (OP2). Surprisingly, EsAureo5 does not contain bZIP domain – instead a LOV with huge N-terminal extension is present. Therefore, as a representative of Aureo5, we chose SjAureo5. In SjAureo5, His153 (NE2) and Lys161 (NZ) interact respectively with OP2 of dG’6 and dG’9. The basic region Arginines, Arg162, Arg151and Arg158 interact with dC’11 (OP2), dT’13 (OP2) and dA’10 (OP2) respectively.

<table>
<tr><th>Name of protein</th><th>Chain No.</th><th>Basic region residues</th><th>Polar interaction details</th><th>Chain No.</th><th>DNA bases</th></tr>
<tr><td rowspan="4">EsAureo1</td><td rowspan="2">C</td><td rowspan="2">RRERNREHAKRSRVRKK</td><td rowspan="4">ARG’355.C /NH2 -- DC ’14.B /OP2<br>ARG’360.C/ NH1 – DG’12.B/ OP1<br>LYS’371.D / NZ -- DA’10.A /OP2<br>LYS’371.C / NZ -- DA’10.B /O5<br>HIS’362 .C / CD2 -- DT’8.A / O4<br>ASN’359.D/ ND2 -- DC’14.A/ N4<br>HIS`362.D /NE2 -- DT`8.B /C6</td><td>A</td><td>CCTTGGCTGACGTCAGCCAAG</td></tr>
<tr><td>B</td><td>CCTTGGCTGACGTCAGCCAAG</td></tr>
<tr><td rowspan="2">D</td><td rowspan="2">RRERNREHAKRSRVRKK</td><td>A</td><td>CCTTGGCTGACGTCAGCCAAG</td></tr>
<tr><td>B</td><td>CCTTGGCTGACGTCAGCCAAG</td></tr>
<tr><td rowspan="4">EsAureo2</td><td rowspan="2">C</td><td rowspan="2">KKRRERNRVLARRTRLRKK</td><td rowspan="4">ARG’190.D/ NH1 -- DC’14.B / OP2<br>ARG’199.D / NH1 -- DG’12.B / OP1<br>ASN’194.C/ OD1 -- DC’14.A /N4<br>ASN’194.C/ ND-- DT’8.B /O4<br>ARG’193.C / NH2 – DG’5.B /OP2</td><td>A</td><td>CCTTGGCTGACGTCAGCCAAG</td></tr>
<tr><td>B</td><td>CCTTGGCTGACGTCAGCCAAG</td></tr>
<tr><td rowspan="2">D</td><td rowspan="2">KKRRERNRVLARRTRLRKK</td><td>A</td><td>CCTTGGCTGACGTCAGCCAAG</td></tr>
<tr><td>B</td><td>CCTTGGCTGACGTCAGCCAAG</td></tr>
<tr><td rowspan="4">EsAureo3</td><td rowspan="2">C</td><td rowspan="2">RRERNREHAKRSRVRKK</td><td rowspan="4">ARG’47.C /NH1 -- DC’7. B /OP2<br>ASN’48.C/ ND2 -- DT’13.A/O4<br>SER’55.C /OG -- DA’10.B /N7<br>ARG’54.C /NE -- DG’9. B /OP2<br>ARG’56.C /NH2 -- DG’9.A /N7<br>HIS’51.D / CE1 -- DG’6.A / C8<br>ASN'48.D / OD1 -- DA'15.B / N7</td><td>A</td><td>CCTTGGCTGACGTCAGCCAAG</td></tr>
<tr><td>B</td><td>CCTTGGCTGACGTCAGCCAAG</td></tr>
<tr><td rowspan="2">D</td><td rowspan="2">RRERNREHAKRSRVRKK</td><td>A</td><td>CCTTGGCTGACGTCAGCCAAG</td></tr>
<tr><td>B</td><td>CCTTGGCTGACGTCAGCCAAG</td></tr>
<tr><td rowspan="4">EsAureo4</td><td rowspan="2">C</td><td rowspan="2">RRERNREHAKRSRIRKK</td><td rowspan="4">ARG’55.C /NH2 -- DG’9.B / OP2<br>ASN’47.D / ND2 -- DG’16.A/ O6<br>ARG’53.D/NH1 -- DG’6.B /OP2<br>ARG’46.C / NH1 -- DC’7.A / OP2<br>HIS’50.D / NE2 -- DG’5.B / C3’<br>HIS’50.C /NE2 --DT’8.A / OP2</td><td>A</td><td>CCTTGGCTGACGTCAGCCAAG</td></tr>
<tr><td>B</td><td>CCTTGGCTGACGTCAGCCAAG</td></tr>
<tr><td rowspan="2">D</td><td rowspan="2">RRERNREHAKRSRIRKK</td><td>A</td><td>CCTTGGCTGACGTCAGCCAAG</td></tr>
<tr><td>B</td><td>CCTTGGCTGACGTCAGCCAAG</td></tr>
<tr><td rowspan="4">SjAureo5</td><td rowspan="2">C</td><td rowspan="2">KRRRNREHAKRSRLRKRVR</td><td rowspan="4">LYS’161.C/ NZ -- DG’9.A / OP2<br>ARG’151.D /NE -- DT’13.A /OP2<br>ARG’158.D/NH2 -- DA’10.A / OP2<br>ARG’158.C /NH2 -- DG’12.B / OP2<br>HIS’153.D /NE2 -- DG’6.B / OP2<br>ARG’ 162.C /NH1 -- DC ’11.B/ OP2</td><td>A</td><td>CCTTGGCTGACGTCAGCCAAG</td></tr>
<tr><td>B</td><td>CCTTGGCTGACGTCAGCCAAG</td></tr>
<tr><td rowspan="2">D</td><td rowspan="2">KRRRNREHAKRSRLRKRVR</td><td>A</td><td>CCTTGGCTGACGTCAGCCAAG</td></tr>
<tr><td>B</td><td>CCTTGGCTGACGTCAGCCAAG</td></tr>
</table>

#### SI-7. : Photoreceptors reported in different algal groups originated through endosymbiosis

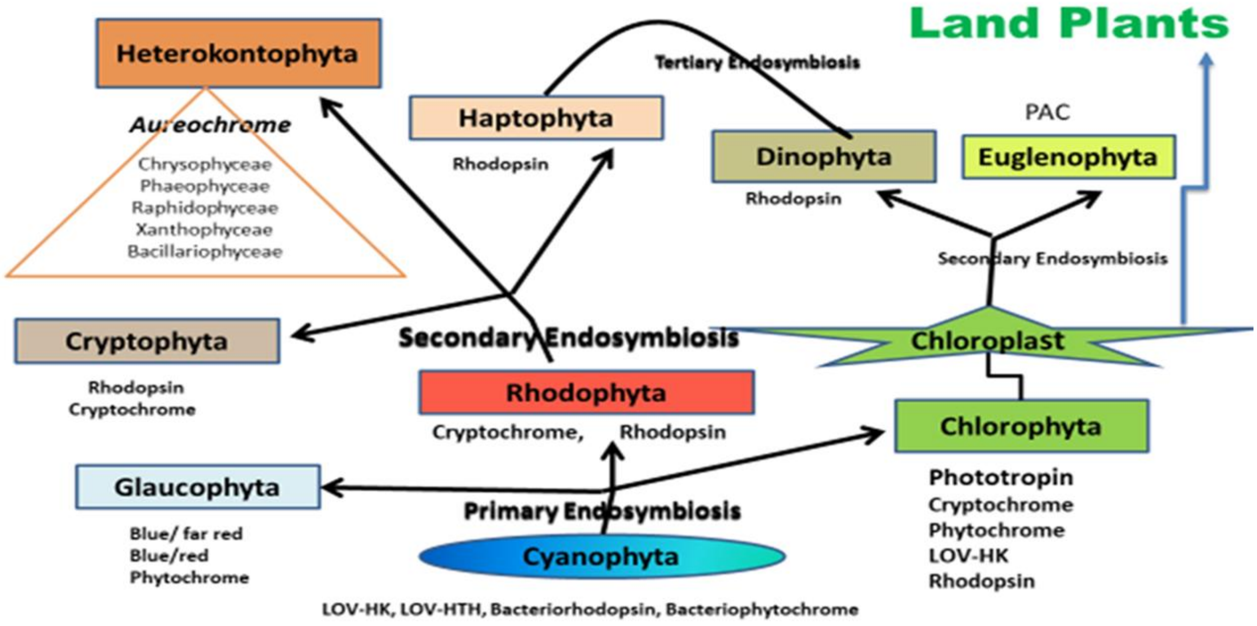


#### SI-8: Overall hydrogen bonding pattern of aureochrome-bZIP protein-DNA

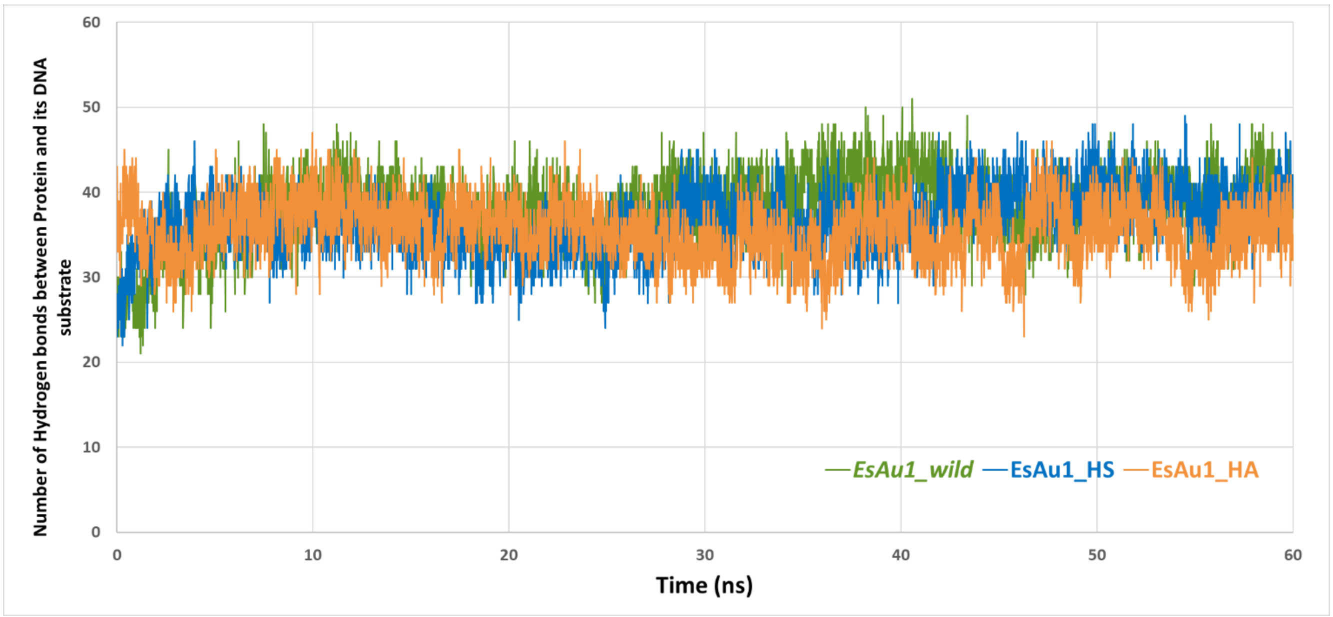
